# *Zymoseptoria tritici* show local differences in within-field diversity and effector variation

**DOI:** 10.1101/2025.01.03.631207

**Authors:** Andrea Tobian Herreno, Pu Huang, Isabella Siepe, Remco Stam

## Abstract

*Zymoseptoria tritici* is a cosmopolitan hemibiotrophic wheat pathogen with a high mutation rate and a mixed reproduction system, leading to challenges in traditional farming management. For successful integration of pest management, especially for early diagnostics of new aggressive or fungicide-resistant lineages, it is critical to understand population diversity in the field. We look at whole-genome sequence data for three datasets to differentiate within field diversity in fields of similar size: one dataset from a newly sampled field population from the United Kingdom and two publicly available datasets from fields from the United States and Switzerland.

Inspection of population structure and diversity features, such as minor allele frequency distribution and clonality, show no within-field structure, the most abundant SNPs are present in low frequency, and European fields have higher clonality. Knowing that effectors play particularly important roles in (a)virulence, we specifically assess effector diversity characteristics. Whereas on a whole-genome scale, we can see separation of the populations at the regional scale, we do not find such separation for the effectors. Moreover, we find that multiple effector haplotypes can be found interspersed within the field and even occur within what has been considered clonal isolates or isolates from a single lesion.

Our analyses highlight that within-field *Z. tritici* diversity is higher than previously reported. Our finding that multiple effector haplotypes can be found within a single lesion might explain the large resistance gene-breaking potential of *Z. tritici*.

## Introduction

*Zymoseptoria tritici (Zt)* is a major wheat hemibiotroph pathogen in the division Ascomycota and the causal agent of Septoria tritici blotch (STB). During the growing season, it reproduces mostly asexually via pycnidia formation after the symptomatic phase has begun. At the end of the season, there is evidence of sexual reproduction where there is pseudothecia formation (Amezrou et al., 2024; Brown et al., 2015; Fagundes et al., 2020; Feurtey et al., 2023; Hartmann et al., 2021; Kellner et al., 2014; Singh, Karisto, et al., 2021). The pathogen’s life cycle with the mixed reproduction system represents a challenge as there is an increase in diversity arising each growth season, combined with resistance structures like chlamydospores and ascospores travelling long distances with the wind (Brunner et al., 2013; Dutta, Croll, et al., 2021; Morais et al., 2017; Rudd et al., 2015; Singh, Karisto, et al., 2021). All these factors combined cause *Zt* to present challenges in terms of management strategies and costs in terms of inoculum diminishing and management of the outbreak, as the symptoms are shown after the colonisation has taken place, and most likely, the inoculum is present in most of the field or on soil deposits. Moreover, the pathogen is rapidly overcoming most commonly used fungicides (Birr et al., 2021; Dutta, Croll, et al., 2021; Feurtey et al., 2023; Klink et al., 2022; Prahl et al., 2023; Stewart et al., 2018).

To accurately manage an outbreak of STB, a sampling strategy with sufficiently deep collection, in the order of hundreds of individuals, is ideal, as it is a more accurate representation of the population causing the disease. This sampling size allows for maximising the diversity captured in sexual pathogens. To maximise the diversity capture, hierarchical sampling is ideal as it can implement an estimation of minimal dispersal distance in the order of tens of cm to long-distance dispersal, potentially minimizing clonal isolate isolation.

Previous studies have also found that there are non-clonal isolates present in the same lesion and leaf layer, remaking the relevance of sampling deeply enough and including different hierarchical distances for Zt as there is an effect of sexual spores in the disease spread, which can be related to the pathogen virulence or the effect of mixed infections within the field (Karisto et al., 2022; B. A. McDonald, 1997; Orellana-Torrejon et al., 2022; Orellana-Torrejon et al., 2022; Siah et al., 2018).

*Z. tritici* has a genome of 39.73 Mbp in 21 chromosomes, of which 13 are considered core and 8 accessory chromosomes (B. A. McDonald et al., 2022; Plissonneau et al., 2018). The dispensable chromosomes also have shown presence-absence polymorphism in a single growing season at the field level, losing chromosomes in a single generation when subjected to stressors. The *Zt* genome contains 531 predicted effector-coding genes (Lapalu et al., 2023). These genes are deemed critical for infection as they are, for example, critical for colonisation through the stomata such as *avrStb6*. Others can modulate and maintain the symptomless phase of the disease in the biotrophic phase like *Mg3LysM* (Alassimone et al., 2024; Mirzadi Gohari et al., 2015; Rudd et al., 2015; Thynne et al., 2024; Yang et al., 2015). *AvrStb9* is another effector-gene that is highly expressed during the asymptomatic phase and is thought to be relevant for the switch to necrotrophy. It codes for a relatively big protein (727 aa) with an S41 protease domain; interestingly, it shows a high variance in its alleles, including deletions in the coding sequence and non-synonymous substitutions, amounting to 44 alleles just in this French isolate collection. The allele variance was key for allowing host recognition evasion (Amezrou et al., 2023). In the necrotrophic phase of the disease, they can be key factors for activating programmed cell death like *Mycgr3_96951* or *Mycgr3_111505* (Thynne et al., 2024). Nonetheless, effectors can also be involved in quantitative responses in their host. Avr3D1 triggers a quantitative response in the cultivars containing Stb7 or Stb12, when recognized the strains containing the avr alleles have reduced colonisation speed and reduced pycnidia formation. This effector also shows high levels of polymorphism due to the high frequency of transposable element (TEs) insertions and point mutations (Meile et al., 2018, 2023).

To date, 23 different *Z. tritici* resistance (R) genes have been mapped, capable of recognizing a wide variety of effectors transcriptionally programmed during both phases of the disease; however, only two of these genes, Stb6 and Stb16q, have been cloned so far. Understanding how pathogen populations evade host recognition, leading to the breakdown of resistance, is crucial for effective disease management (Amezrou et al., 2023; Brown et al., 2015; Kettles & Kanyuka, 2016; Rudd et al., 2015). Thus, understanding genetic variation in *Z. tritici* is essential for gaining insights into the pathogen’s evolution, particularly regarding the rise of fungicide resistance and assessing the potential durability of deployed R genes. The pathogen’s large effective population size (N_e_) enables the emergence of fungicide resistance mutations driven by continuous fungicide exposure. In addition, cultivar (R gene) selection creates selective pressures that lead to a vast array of unique genotypes. These genotypes, often possessing enhanced fitness in terms of aggressiveness or overall resilience, can contribute to the devastating outcomes of STB outbreaks (Birr et al., 2021; Croll & McDonald, 2017; Garnault et al., 2021; Hartmann et al., 2018, 2021; Hellin et al., 2021; Kiiker et al., 2021; Klink et al., 2022; Lendenmann et al., 2015; M. C. McDonald et al., 2019; Vestergård et al., 2023).

The accessory genome in *Zt* contributes to the virulence of the strain based on presence- absence variation, and some accessory chromosomes can recombine with core chromosomes, creating a new source of diversity for the pathogen (Schotanus et al., 2015). Yet, it should be noted that only 11 effectors have been annotated on the dispensable chromosomes. There are also other means by which *Zt* generated genomic diversity. There is an abundance of TEs in the genome, allowing the translocation of various effector genes (Lorrain et al., 2021; Oggenfuss et al., 2021; Singh, Badet, et al., 2021). Another source of variation via repeat-induced point mutations (RIPs) also contributes to the high mutation rate (μ) evidenced in the literature (Hartmann et al., 2021; Singh et al., 2022; Van Wyk et al., 2021).

On the global level using core genome SNP diversity data, *Zt* shows well-supported clades with a single genetic cluster in Europe, two in North America, one in South America, one in Australia, and another in New Zealand. Genetic exchange exists between the Americas and Europe and between Oceania and Europe. However, the hotspots of diversity are the clades in northern Africa and the Middle East, wheat’s origin, and domestication sites (Feurtey et al., 2023). However, studies using SSR markers denote a smaller-scale population structure confined to the regional level in Tunisia and also in France where diversity is bigger compared to other countries in Europe, in both studies, they remark on the contribution that sexual reproduction has to the overall diversity denoting the varied combination of haplotypes, while still observing private alleles at the SSR level (Chedli et al., 2022; Siah et al., 2018).

These unique genotypes can have varying degrees of fitness as they can affect fitness terms of virulence that award them not only differential infection frequency and dispersal, but also factors like speed of disease progression and immune response modulation, conferring an adaptive advantage for these effector haplotype configurations depending on the host context or affecting the survival in the case of fungicide exposure. There is evidence of host specialisation in Zt at the transcriptional level and also SNPs associated with various cultivars (Amezrou et al., 2024; Hartmann et al., 2017; Karisto et al., 2022; Kellner et al., 2014; Lorrain et al., 2024; Orellana-Torrejon et al., 2022; Suffert et al., 2015). Understanding how these haplotypes are distributed in the fields can give insight into effector allele dominance within and between fields.

In-depth population structure analysis at the field level is limited to date, but can shed light on the population dynamics behind the variability contained on the field. As non-clonal isolates, and also a mixture of virulent and avirulent isolates can be contained inside a single lesion, understanding how clonality spreads during the disease outbreak can be a key factor in designing management strategies (Bernasconi et al., 2022; Karisto et al., 2022; Orellana-Torrejon et al., 2022; Siah et al., 2018). Population genomics analyses can be biased. Depending on the research question, different criteria for data filtering and downstream analyses might need to be applied (Everhart et al., 2021). Previous studies of within-field diversity focused on general diversity statistics and unique isolates; for this purpose, the sequencing data was filtered to maximize variant call confidence and alleles of intermediate frequency by using minor allele frequency filtering and SNP pruning. This data mining is well suited to finding differences at the continental or regional level and estimating ancestral population size (Feurtey et al., 2023; Hartmann et al., 2018; Singh, Karisto, et al., 2021). We want to specifically assess the overall within field diversity and the putative effects thereof on putative virulence or avirulence factors such as effectors. To this effect, we analyse three different field data sets to contrast different filtering settings and apply more inclusive analyses to assess the genetic structure of this pathogen in the field regarding clonality, minor allele frequency distribution, and nucleotide diversity and specifically look at effectors as putative virulence or avirulence factors.

## Materials and Methods

### Data set sources

Sequencing data for the Swiss and US fields were sourced from Feurtey et al., 2023. ADAS collected the field samples from the UK at their Rosemaund field station (Hereford, UK). Sequencing was done by Microsynth and sequence data was deposited under ENA Bioproject PRJEB81422. The sequencing data for the Swiss and American fields is available in NCBI in the BioProject PRJNA596434. The GPS coordinates for each isolate are present in the supplementary material (Table S1). Selection of samples from UK field was made by human choice, aiming to represent different areas of the fields and having a wide range of distances between samples, going from clusters only several meters apart to single samples more than a hundred meters separated from each other, the collection was made in 2021 at Growth Stage 83. The field contained a single wheat cultivar Graham. The Swiss field had a three-time-point collection strategy in twelve cultivars and subsequent fungicide treatments; the material was collected from different plants in the same cultivar plots (details see: Singh, Karisto, et al., 2021). The US field-collected isolates have been sampled over a larger geographical distance than the other fields. Samples were obtained from two cultivars; Stephens and Madsen (Zhan et al., 2005).

### Depth assessment

We aligned the reads to the reference IPO323 using BWA v.0.7.17 with the option bwa mem. The resulting Sam files were sorted and indexed using SAMtools v.1.17 (Li et al., 2009). The quality check for the Bam files was done using the Picard dependencies from GATK v.4.0.5.1 with the option ValidateSamFile (Poplin et al., 2017). Then we checked the depth of the dispensable chromosomes using samtools --depth and ranked them by thresholding, depth < 0.25 absent, 0.25 > depth < 0.625 partial deletion, 0.625 > depth < 1.5 presence and depth >1.5 duplication (as defined in Singh, Karisto, et al., 2021).

### Alignment and variant calling

After Bam file validation, we did the multi-sample variant calling with GATK HaplotypeCaller for the subset of isolates per field and GenotypeGVCF with a maximum of 2 alternate alleles and 6 maximum genotypes per site. Then we used quality filters with the VariantFiltration tool with the following cut-offs: MQ < 20.0, QD < 2.0, QUAL < 30.0, SOR > 3.0, FS > 60.0, DP < 10, -2.0 > ReadPosRankSum < 2.0, -2.0> MQRankSum < 2.0. Using SelectVariants, we only kept SNPs and allowed for only 2 alleles and a maximum of 6 genotypes.

For the strict filtering (described in Singh, Karisto, et al., 2021), we hard-filtered our variant call further using VCFtools v.0.1.16 for a per locus genotyping rate > 80% with the option -- max-missing 0.8 and kept a Minor Allele Count (mac) of 1, with the option --mac 1. For our custom (relaxed) filtering we kept the settings for the GATK pipeline but only applied the minor allele count filter –mac 1 with vcftools. The final genotyping rate was extracted with plink 1.90b6.21 using the --missing option with the bed file corresponding to each field data set. We extracted all the minor allele frequency (MAF) distributions using the option –freq. We made all figures with ggplot v.3.5.0 in R v.4.2.1 (2023).

### Clonal quantification in the fields

Using the data for the SNP calling, we used R v.4.2.1 with the package poppr v2.9.5 (Kamvar et al., 2014) to determine the genetic distance using poppr::bitwise.dist (Hamming distance, Hamming relative distance). With this distance matrix, we used the hclust function from the base and the clonal distance defined by Singh, Karisto, et al., 2021 of distance threshold = 0.01, representing approximately 0.01% of the total SNP count, equivalent to an average of 132 SNPs. We plotted a Neighbour-Joining tree (NJT) with the defined clusters, then used tidyverse v.2.0.0 to extract and count samples clustered together whose clade was shorter than 0.01 in subsets for isolates coming from the same plant or plot, same leaf, and same field (Wickham et al., 2019).

### Admixture analysis

The admixture analysis was done with the pruned high-confidence-GATK filtered variant call using plink --indep-pairwise using a 50 SNPs window, shifting every 10 SNPs, with 0.2 variance inflection threshold. We applied a minor allele frequency (MAF) filter 0.05 to this pruned data and ran admixture v1.3.0 with K1:10. The results were plotted using ggplot v.3.5.0 in R v.4.2.1.

### Calculation of invariable regions

For the calculation of invariable regions, we used the definition of telomeric and centromeric regions in *Z. tritici* determined in Schotanus et al., 2015. We define as invariable regions, stretches of the chromosome that have no SNPs called with the reference IPO323. We used Bedtools v2.31.1 with the option –intersect and the reference annotation to IPO323 of contig length for all the chromosomes to determine the uncorrected invariable regions in the whole genome of each field (Quinlan & Hall, 2010). To correct the regions to exclude centromeres and telomeres, we used a custom Python script that flagged the invariable regions, telomeres, and centromeres. The genome was complemented with effectors annotation from the latest available IPO323 annotation (Lapalu et al., 2023, p. 202). This script can be found at https://github.com/PHYTOPatCAU/ZymoFieldDiv.

### Population genetics statistics

We performed a subsetting analysis to estimate the effect of sample size by creating sets of randomly selected isolates into groups outputted into a text file with the shuf function in a python base environment and then counting the number of SNPs as given by VCFtools v.0.1.16 with the --keep option and --mac 1 to keep the polymorphic sites(Danecek et al., 2011). To assess which growth model was best fitted to our data, we used the r4pde, minpack.lm and the knittr in R (Del Ponte, E. M, 2023; Elzhov Timur et al., 2023; Xie Y, 2024). We calculated the residuals and R^2^ of the logistic, exponential, power, and linear models to determine which had the higher explanatory power.

Each country was fitted with a power growth model in R v.4.2.1 using the package minpack.lm v1.2.4, with the nlsLM function (Elzhov Timur et al., 2023). The flattening point for each curve was calculated using the fitted curve, assuming the plateau was when the derivate was smaller than 1^-6^ for each combination of Country-Filter combination Because of the slow flattening of the original analysis, we also included a threshold where the plateau would be reached if the slope corresponded to an increase of less than 1% of the total number of SNPs found in each Country-Filter combination.

We calculated the Principal Component Analysis (PCA) of the three fields, using R with the package SNPrelate v.3.13 to input the vcf file and convert it into an SNP GDS format. Then, we calculated the PCA with the snpgdsPCA function and plotted the first two eigenvalues (Zheng et al., 2012).

We calculated the geographic distance matrix using the GPS coordinates for the isolates in UK field using geosphere v1.5-18 after we paired the geographic distance matrix and the genetic distance matrix calculated with poppr to assess the relation between both (Hijmans, 2022). The relation was calculated using the R base function mantel.

The diversity statistics of nucleotide diversity (pi) and Tajima’s D were calculated using the scikit-allele v1.3.7 toolset (Alistair Miles et al., 2024); both statistics were calculated using the entire chromosome variant call using a custom python script found at https://github.com/PHYTOPatCAU/ZymoFieldDiv. The figures were plotted in R v.4.2.1 with ggplot v.3.5.0 (R Core Team, 2023).

We obtained the Effectorome annotation from the latest available annotation in JGI for Zt (Lapalu et al., 2023) and used it to extract only the SNPs in effectors for each field and did a PCA analysis with this dataset. Furthermore, we also analysed the haplotype variation of 9 functionally characterized effectors using minimum spanning networks generated using poppr in R, including *avrStb6, avrstb9 and avr3d1* (Amezrou et al., 2023; Lapalu et al., 2023; Rudd et al., 2015). We also describe the distribution for the haplotype of *avrStb6, avrstb9 and avr3d1* (*ZtIPO323_060700*), *ZtIPO323_119610* and *ZtIPO323_117500* across the UK field using the GPS coordinates for each isolate.

## Results

### The genotyping rate for the three fields is comparable, yet SNP numbers vary

We compare *Z. tritici* genomic diversity data from three different wheat field sites. A newly sampled field from the UK and two previously published fields in Switzerland and the US (Singh, Karisto, et al., 2021). The datasets had different average depths as they were different sequencing efforts; overall, we have a coverage of more than 30x across all of them, denoting the deep sequencing needed for the variant calling analyses. The Swiss dataset has an average coverage of 69x, the UK has coverage of 47x, and the US has a coverage of 32x. With the strict filtering done in Singh, Karisto, et al., 2021, we detected 1,438,066 SNPs in the Swiss field, a similar number of SNPs as in the previous study using a subset of 158 isolates. In the UK field, we detected 1,193,611 SNPs from 160 isolates. The US fields deviate as they have a smaller sample size of 97 isolates. We detected 572,804 SNPs for the US field. We show an average genotyping rate of 97.09% for Switzerland, 96.73% for the UK, and 96.86% for the US (Figure 1. a, Figure 2. a, FigureS1).

**Figure 1.**
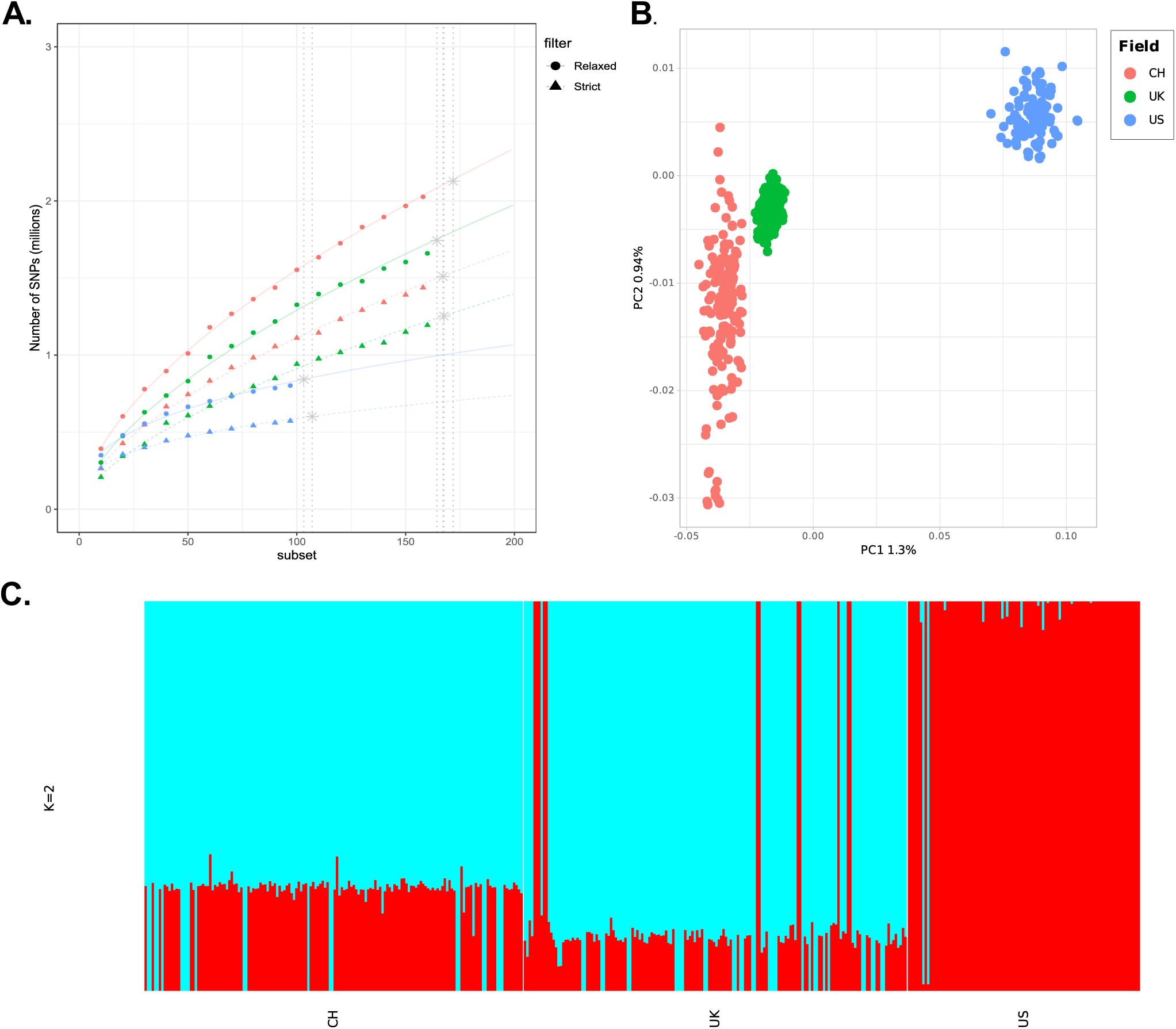
Between fields overview. **a.** Number of segregating SNPs for the relaxed filter dataset, calculated using VCFtools 0.1.16. the dashed and horizontal lines represent the predicted growth fitted to the power model, and the vertical lines correspond to the predicted sampling fraction where the curve flattens, assuming a threshold of 1% increase in the total SNP count. The grey stars represent the predicted maximum SNPs at the adjusted plateau**. b**. PCA with the whole genome SNPs for each field w/o outliers calculated using SNPRelate::snpgdsPCA. **c.** Admixture analysis from the three fields using the relaxed filtering from the core genome with MAF 0.05 using Plink --maf 0.05.

**Figure 2.**
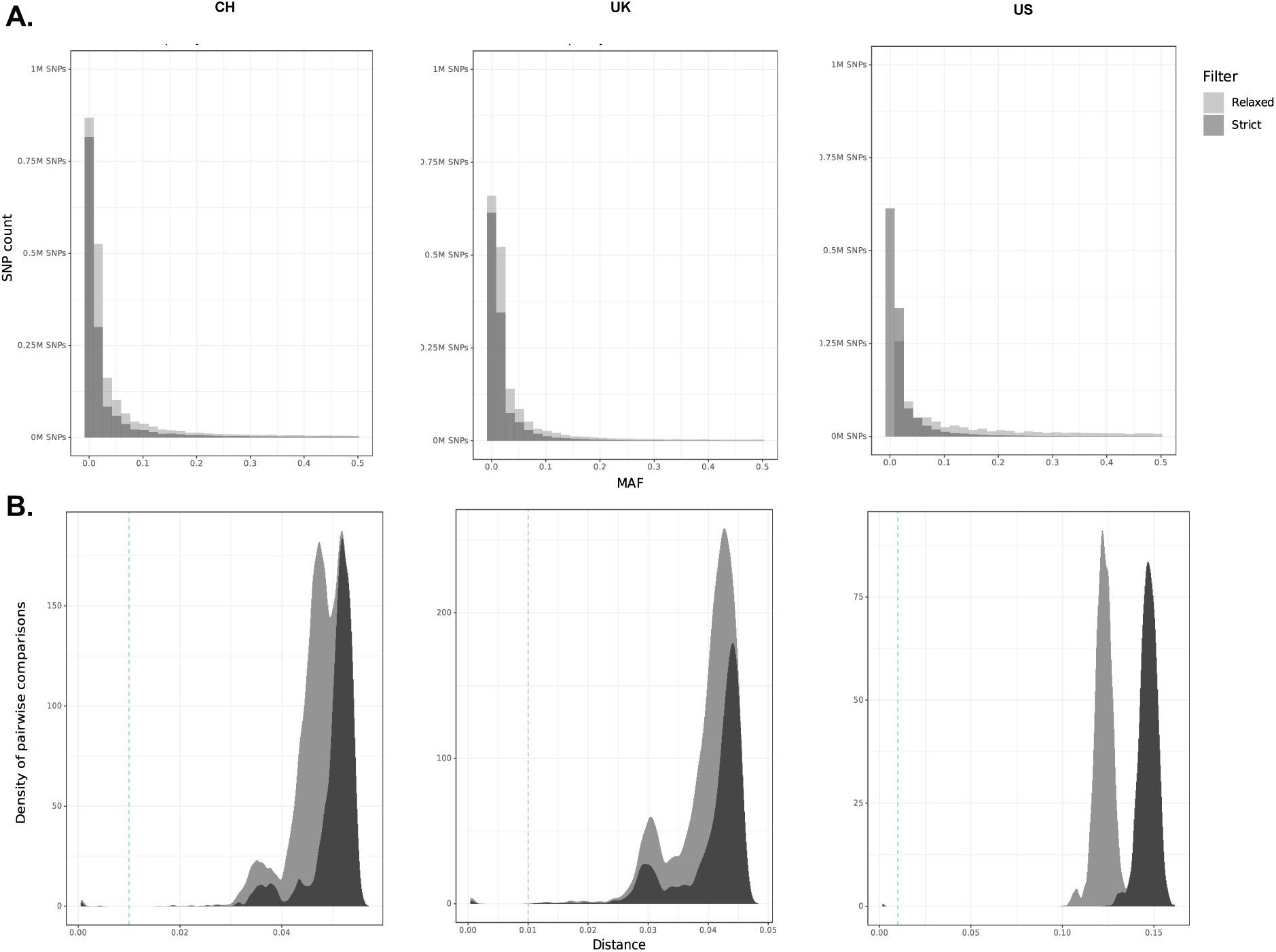
Minor allele frequency distribution of the three fields and histograms of the genetic distance of the Strict and Relaxed filtered datasets. The Minor Allele Frequency (MAF) distribution was calculated for each field using Plink --freq. The genetic distance of the core genome was calculated using Poppr in R with the option bitwise.dist. **a.** MAF distribution per field, MAF for the Strict filtering, the total SNP counts are CH 1,438,066 SNPs, the UK 1,193,611 SNPs, and the US 572,804 SNPs. For the Relaxed filtering, the SNP counts are CH 2,026,783 SNPs, UK 1,660,037, and US 802,498 SNPs. **b.** Histograms of within-field genetic distance normalised by count of pairs of comparisons.

We repeated the analysis keeping only the minor allele count filter (mac 1 option) from the downstream processing with vcftools from the variant call pipeline reported in the Swiss field survey (Singh, Karisto, et al., 2021) so we could exclude false positive calls for all fields, whilst at the same time take a more lenient approach where we keep all SNP calls present in the accessory chromosomes and SNP with partly missing data in some isolates. We keep all up to 6 possible haplotypes, and not only biallelic calls. With this relaxed approach, we detected 2,026,783 SNPs in the Swiss field, 1,660,0377 in the UK field, and 802,498 in the US field (Figure 1. a, Figure 2. a). Thus, we recovered significantly higher amounts of SNPs in all the data sets. We show an average genotyping rate of 87.04% for the UK, 86.04% for Switzerland, and 84.25% for the US (Figure S1).

### The saturation curves for the number of SNPs given a subsampling fraction are similar between populations

The Swiss field has the highest number of SNPs, followed by the UK, and the US has the fewest SNPs per subset. We tested different models on the SNP rarefaction curves to assess whether our sample sets capture the full extent of the within-field variation (Table S2). The power model showed the best fit. Thus, we modelled the predicted plateau for each combination of country-field by fitting the existing data to a power growth model curve (Figure 2. d). Assuming that the diversity bulk will be collected until we only observe an increase of 1% in SNP count, the predicted number of samples necessary for reaching the plateau varies for the Swiss, UK and US fields. The difference between the filtering is small. in the Swiss field, we need 168 samples to reach the plateau with the strict filters and 171 with the relaxed filters. The UK field reaches the plateau with 166 isolates for the relaxed filter and 161 isolates for the strict filtering. The US field shows a predicted inflexion point at 83 or 84 isolates for the strict and the relaxed filters, respectively (Table 1). These plateau predictions show that we capture the most relevant variation in the three fields and confirm that the US field shows significantly less genetic diversity than those in the UK and Switzerland.

**Table 1.** Modelling results for sampling efforts. The curve results are based on the SNP saturation curve and fit a logistic growth model.

| Country | Filter | SNPs at plateau | Isolates a plateau |
| --- | --- | --- | --- |
| CH | Relaxed | 2,128,122 | 172 |
| UK | Relaxed | 1,743,039 | 164 |
| US | Relaxed | 842,622 | 103 |
| CH | Strict | 1,509,969 | 167 |
| UK | Strict | 1,253,291 | 167 |
| US | Strict | 601,444 | 107 |

### The admixture analysis and PCA analysis shows two clusters separating Europe from***the US***

Admixture analysis and the PCA show two distinct genetic clusters, one from the European and another from the US fields. There is evidence of contribution from the European fields that gave rise to the US field. Some US isolates seem to share the same ancestry as those in the UK (Figure 1. c). The visible outliers in the PCA analysis of the whole genome in the European fields show more diversity in the Swiss field than in the UK. (Figure1.a, Figure1.b, Figure S2. b).

### Adjusted filtering recovers intermediate frequency SNPS

The MAF distribution for the fields shows a skew towards rare alleles, consistent with previous studies, regardless of filtering settings. This means that the bulk of the variation is present at a low frequency in the fields. Our relaxed filtering settings increase the number of SNPs at all allele frequencies and generate a relatively larger increase in intermediate frequency SNPs, indicating that sites that had previously been filtered out contain valuable data that might help describe within-field dynamics (Figure 2a). We see a different magnitude of SNPs detected, probably due to the differences in sample sizes. The increase in intermediate frequency SNPs appears to be larger in the US than in the UK and Swiss fields. Yet, also in the US, we can see the skew towards a large frequency of rare alleles characteristic for the *Zt* populations, indicating that there are no signs of bottlenecks in these fields. This is consistent with previous studies that find no evidence of a recent bottleneck event both at the global and regional scale (Croll & McDonald, 2017; Hartmann et al., 2018; Singh, Karisto, et al., 2021).

### The European fields have less genetic distance than the US field

We calculated pairwise genetic distance between all isolates within the fields. We show high similarity in the pattern of the genetic distance distribution between the fields, where most isolates are equally distant from each other, drawing a peak near the maximum genetic distance, regardless of filtering settings (Figure 2. b). This is consistent with the MAF pattern we saw in Figure 2a. The US has a maximum genetic distance of ∼ twice the size of the maximum found in the European fields. The filtering did not affect the genetic distance distribution between both filtering settings (Table 2). Our threshold for clonality is 1% of the total diversity, i.e., total SNP count per field. We also show the Hamming distance (pairwise SNP dissimilarity) (Table 3, Figure S4). We observe similar numbers for mean distance in the US and CH fields, and a much smaller SNP distance in the UK. Given the smaller total SNP count in the US, we show that the mean average distance in the US field is bigger than the other fields.

**Table 2.** Genetic distance distribution per filtering parameters.

| Field | CH | UK | US | Filtering |
| --- | --- | --- | --- | --- |
| Minimum genetic distance | 0.0007 | 0.0006 | 0.002 | Relaxed |
| Mean genetic distance | 0.045 | 0.039 | 0.116 | Relaxed |
| Maximum genetic distance | 0.053 | 0.046 | 0.128 | Relaxed |
| Minimum genetic distance | 0.00053 | 0.0004 | 0.0021 | Strict |
| Mean genetic distance | 0.05 | 0.041 | 0.116 | Strict |
| Maximum genetic distance | 0.057 | 0.048 | 0.128 | Strict |

**Table 3.** Genetic SNP distance distribution per filtering parameters.

| Field | CH | UK | US | Filtering |
| --- | --- | --- | --- | --- |
| Minimum genetic distance | 1,353 | 860 | 1,672 | Relaxed |
| Mean genetic distance | 84,294 | 59,369 | 94,340 | Relaxed |
| Maximum genetic distance | 98,039 | 70,628 | 103,991 | Relaxed |
| Minimum genetic distance | 773 | 509 | 981 | Strict |
| Mean genetic distance | 72,171 | 48,672 | 83,983 | Strict |
| Maximum genetic distance | 82,018 | 57,647 | 92,430 | Strict |

### Clone probability is similar in the European fields

We calculated the clonal isolates using the clone definition from Singh, Karisto, et al., (2021), being clones sharing < 1% of the total SNP count between a pair of isolates. These clone clusters can be easily identified in Figure 2. b as a small peak left to the clonality threshold in light blue. A few pairwise comparisons are below the clonality threshold in all fields, and filtering settings do not affect the outcome. The European fields have more clones than the US. All three fields showed the presence of clonal isolates (Table 4; Figure 2. b). The US field has the fewest clones, with only 6 isolates clustering below the threshold as three pairs of clones. The CH field showed 28 isolates clustering as clonal, mostly in pairs or groups of three corresponding to the same plots. In the UK, we found 54 isolates below the clonality threshold, where most were grouped as pairs in the same plant or, in two cases, clusters of three isolates in nearby GPS points (Figure S3).

**Table 4.** Clone probability per field. The probability of finding isolates below the genetic distance threshold (<0.01).

| Field | CH | UK | US |
| --- | --- | --- | --- |
| In the same plant/plot | 0.893 | 0.692 | NA |
| In the same field | 0.177 | 0.319 | 0.062 |

***There is presence-absence variation of accessory chromosomes between fields*** Knowing that large-scale presence-absence variation of the accessory or dispensable chromosomes can have large effects on effector virulence, we compare PAV patterns between fields. The US field has very little variation in the dispensable genomes. Most accessory chromosomes are present in all the isolates, partial deletions exist in all chromosomes, and a single duplication event is detected on chromosome 14. The Swiss field shows more variation, with evidence of the absence of chromosomes 15, 17, 19, 20, and 21 and duplications in chromosomes 17, 18, 19, and 21 in up to 3,95% of the isolates. The UK field shows more deletions in all accessory chromosomes, up to 16,9% for chromosome 15 and as low as 3,75 for chromosome 16. This field only presents duplication events in chromosome 14 (Figure 3). No isolate in the three fields shows deletion of all accessory chromosomes. Chromosome 18 shows the lowest coverage in all three fields, but only in the UK field; we see a complete deletion. Also, in the UK field, we see that the most absent is chromosome 16 (Figure 3, Figure S6). This presence-absence variation does not impact effector diversity significantly, as only 11 predicted effectors are found in the dispensable genome (Figure 4).

**Figure 3.**
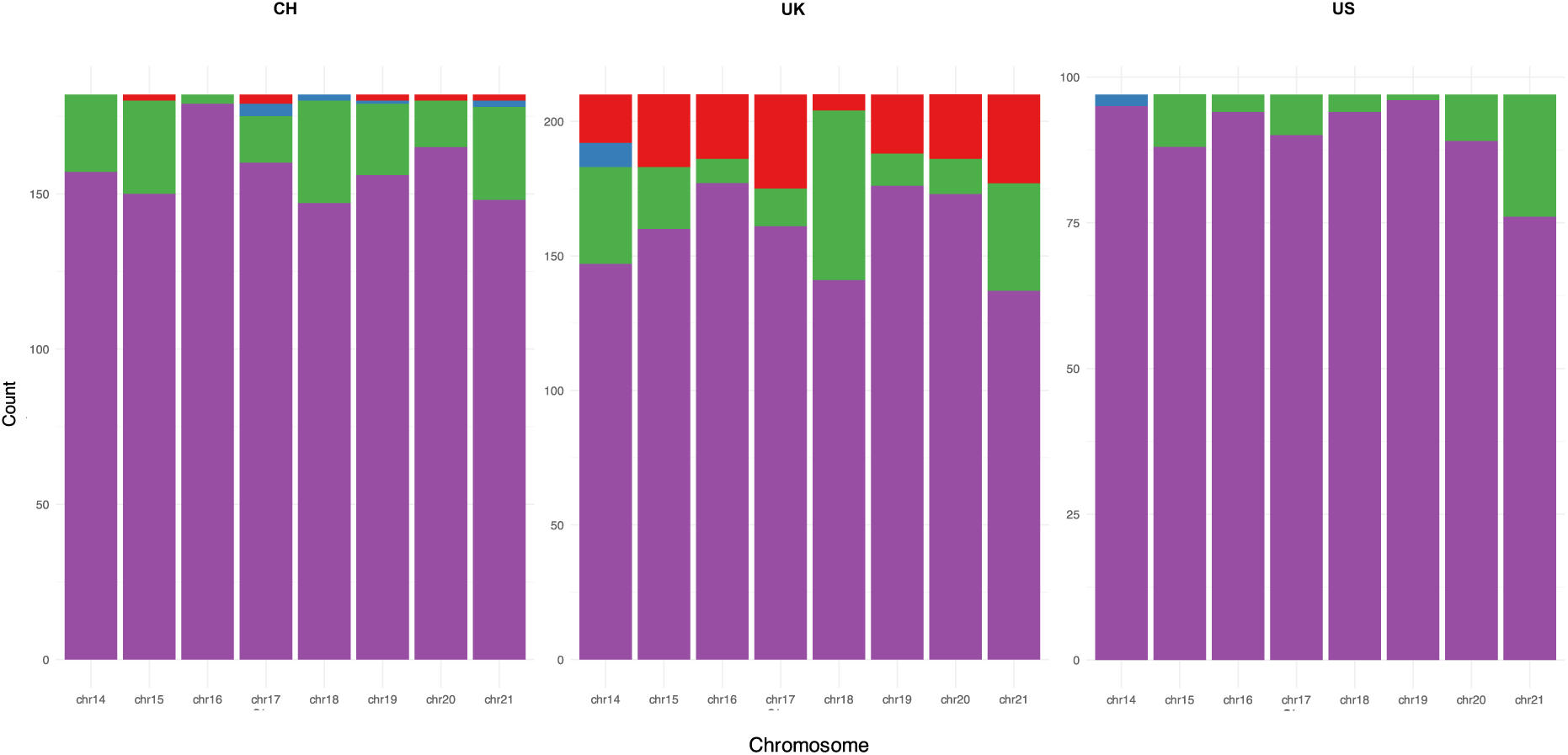
Presence-Absence Variation of the dispensable genome. PAV in the fields As in Singh, Karisto *et al.,* 2021 depth definitions are as follows: depth < 0.25 = absent, 0.25 > depth < 0.625 = partial deletion, 0.625 > depth < 1.5 = presence and depth > 1.5 = duplication.

**Figure 4.**
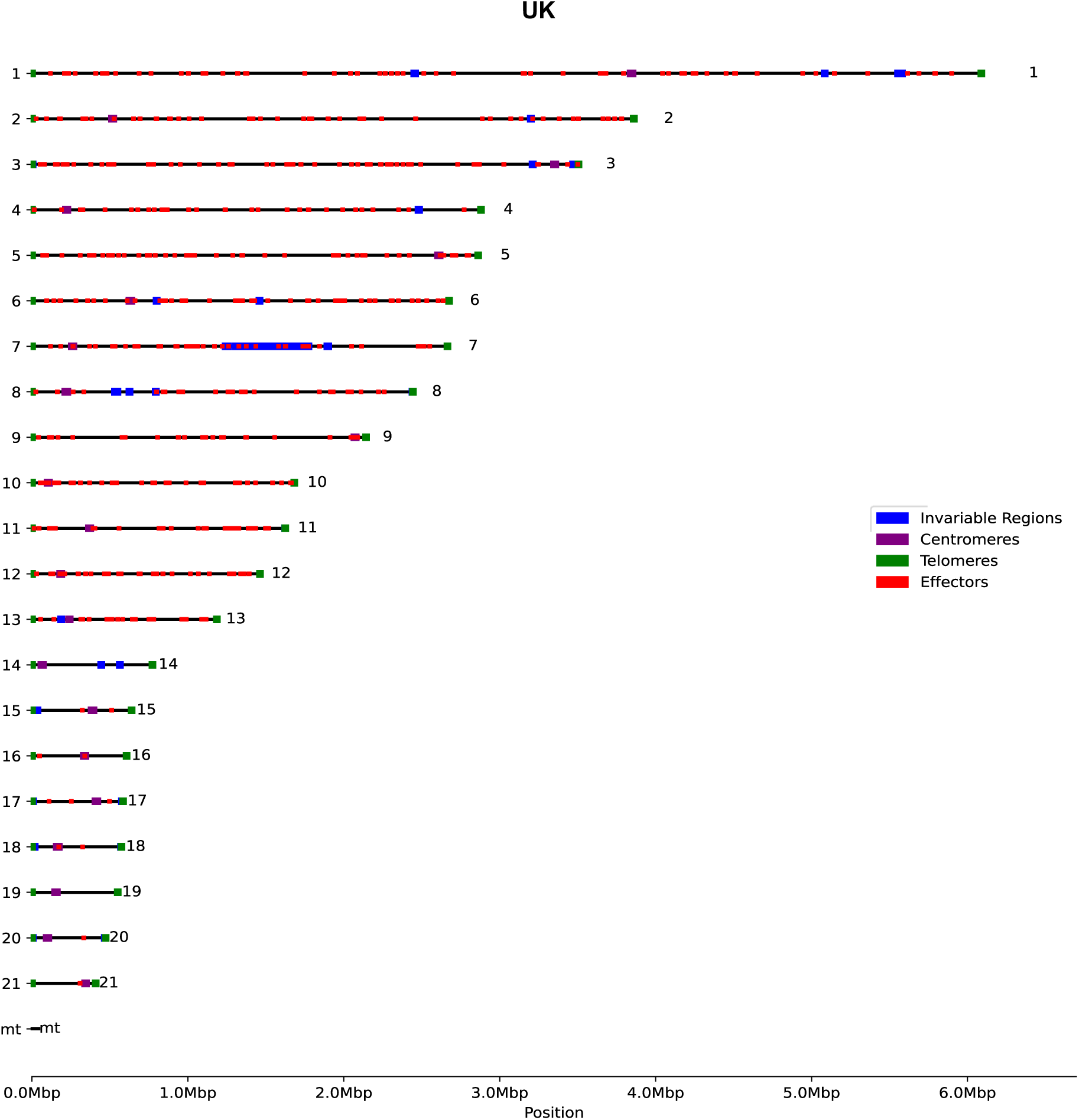
**Invariable regions in the whole genome**. The UK field was screened for invariable regions per chromosome using the reference IPO323 for comparison. The telomeric and centromeric regions were annotated as determined by Schotanus *et al*. 2015. Using a custom script, the invariable regions were corrected using the location for centromeres and telomeres.

### The invariable regions are different within each field

Next, we checked whether the genetic diversity is always similarly distributed across the genome. The invariable regions in each field show a different pattern; the European fields show more similarity with the least number of invariable regions compared to the US field, which has more invariable regions. One feature common to all fields in the invariable region in chromosome 7 is approximately 50kb in length. Only four to seven putative effectors in this region are found in the newest annotation (Lapalu et al., 2023), showing a very small effect on the overall effector diversity (Figure 4, Figure S5).

### The nucleotide diversity in the US is bigger than in the European fields

The SNP count was higher in the European fields than in the US field, regardless of which filtering we used (Figure 5. a). When looking specifically at effector diversity in each field, we find 24,721 SNPs in the Swiss field, 20,066 SNPs in the UK field, and 8,172 SNPs in the US field in the curated effectors set of 526 genes using the relaxed filtering settings.

**Figure 5.**
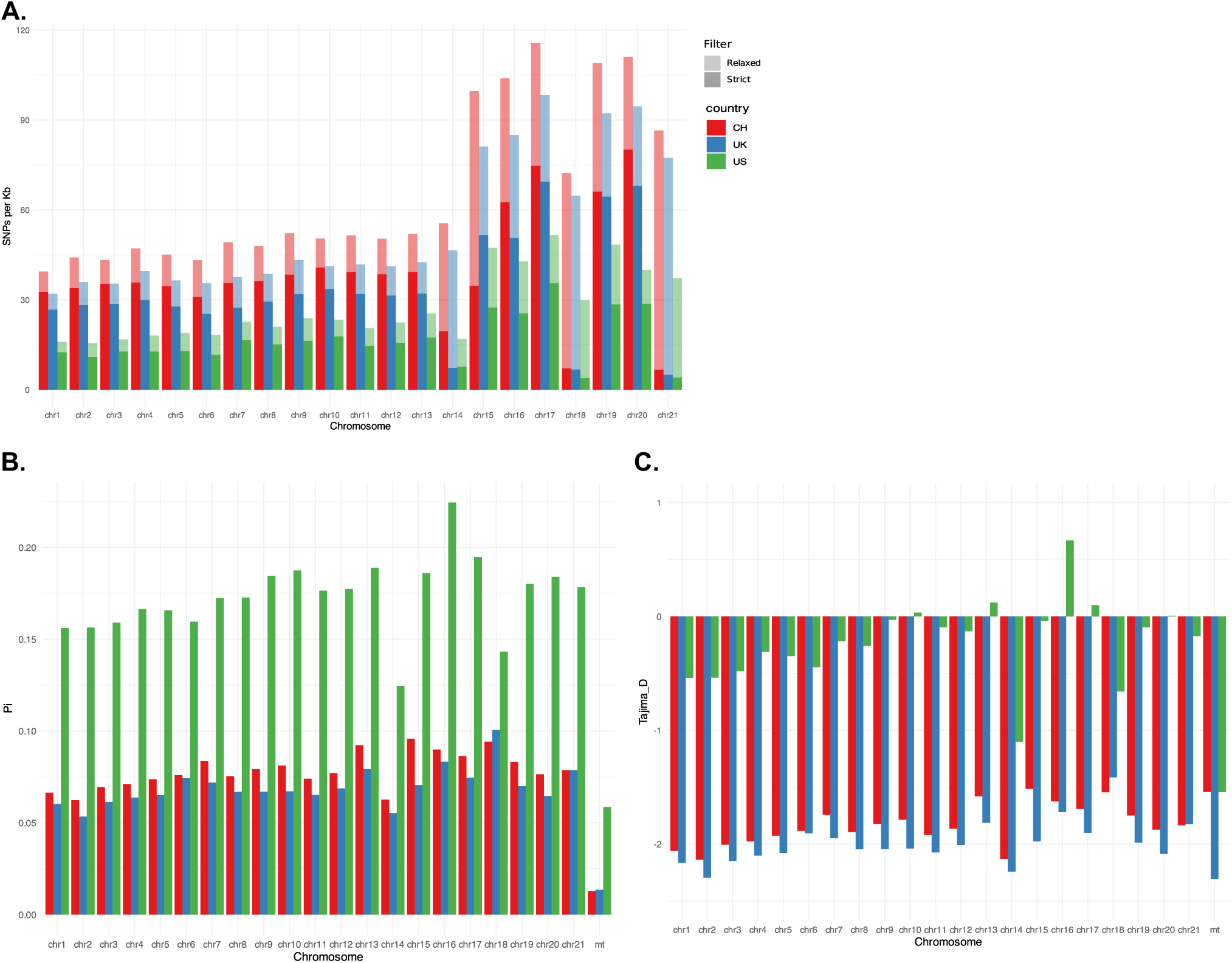
SNP distribution and diversity statistics of the fields. **a.** SNP density measure per chromosome in terms of SNP per Kb, corrected by the chromosome size. **b.** Nucleotide diversity (Pi) per chromosome in the three fields. **c.** Tajima’s D scores per chromosome for the three fields. Nucleotide diversity and Tajima’s D were calculated using scikit-allele v.1.3.7 in a Python 3.9 env.

Interestingly, the overall nucleotide diversity (Pi) was higher in the US field, suggesting fewer shared SNPs within the American population (Figure 5. b). A few non-neutral Tajima’s D scores were negative and found in both European fields on the core chromosomes and chromosome 14. The UK field also shows a significant negative Tajima’s D value in accessory chromosome 20 and the mitochondrial genome (Figure 3. c). This has consequences for the effectors, which are majorly located in the core chromosomes, chromosomes 1 to 13 (Figure 4), causing constraints in haplotypes and overall diversity probably determined by host barriers. For the UK, there is almost always a Tajima’s D index smaller than -2 in the core, chromosomes 14 and 20. In contrast, for the Swiss field, we only see such values in chromosomes 1 to 3 and 14. This might be because the field had multiple host genotypes with variable genetic backgrounds, allowing for multiple haplotypes to be present.

### Effector diversity is similar between the fields

We calculated the nucleotide diversity and Tajima’s D scores for the annotated effectors. We showed that in correspondence to the SNP number per field, the Swiss field has the biggest diversity, followed by the UK and, finally, the US (Figure 5. a). This contrasts with what happens in the chromosome context, where the US has the most nucleotide diversity among the fields. However, when we see the Tajima’s D scores, we observe a general correspondence to the genomic contexts, where most significantly positive Tajima’s D scores are present in the US field. A few Effectors also have high Tajima’s D scores in the Swiss and UK fields. Nonetheless, most effectors remain neutral (between -2 and 2) or have negative scores for all fields. Several effectors sow very low Tajima’s D. (Figure 5. b, Figure 6. a). In Table 5, we show the values of five functionally characterised effectors. Here, we can see that the European fields both have less nucleotide diversity than expected for the effector *ZtIPO323_117500*, the UK even showing a significantly constrained diversity. A remarkable feature is that *avrStb6* has more diversity than expected in the US and not in the European fields. avr3D1 displays neutral statistics in all the fields, but avrStb9 has less diversity than expected in the UK field.

**Figure 6.**
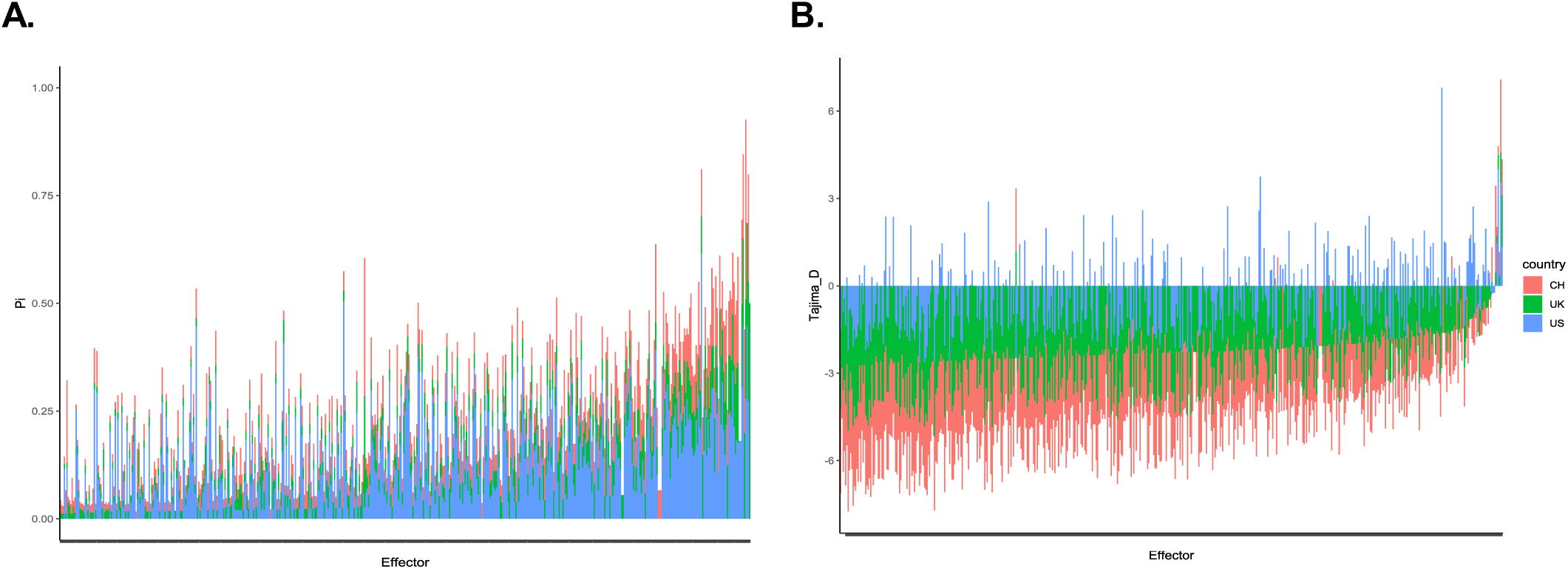
Diversity statistics for the annotated effectors. **a.** Effector nucleotide diversity (Pi) in the fields. **b.** Tajima’s D scores for the Effectors in the Fields. Nucleotide diversity and Tajima’s D were calculated using scikit-allele v.1.3.7 in a Python 3.9 env. The Effector genes annotation and localisation were found in Rudd *et al.,* 2015 and Amezrou *et al.,* 2024.

**Table 5.** Diversity statistics for functionally characterised effectors

| Effector | Pi | Tajima's D | Field |
| --- | --- | --- | --- |
| *avrStb6* (*ZtIPO323_060700*) | 0.068 | -1.946 | CH |
| *avr3D1* | 0.279 | 1.735 | CH |
| *avrStb9* (*ZtIPO323_007790*) | 0.091 | -1.373 | CH |
| *ZtIPO323_117500* | 0.052 | **-2.213** | CH |
| *ZtIPO323_119610* | 0.183 | 0.088 | CH |
| *avrStb6* (*ZtIPO323_060700*) | 0.031 | **-2.359** | UK |
| *avr3D1* | 0.060 | -1.799 | UK |
| *avrStb9* (*ZtIPO323_007790*) | 0.036 | **-2.379** | UK |
| *ZtIPO323_117500* | 0.050 | **-2.241** | UK |
| *ZtIPO323_119610* | 0.098 | -1.132 | UK |
| *avrStb6* (ZtIPO323_060700) | 0.341 | **2.423** | US |
| *avr3D1* | 0.147 | -0.714 | US |
| *avrStb9* (ZtIPO323_007790) | 0.085 | -1.438 | US |
| *ZtIPO323_117500* | 0.192 | -0.031 | US |
| *ZtIPO323_119610* | 0.301 | 1.668 | US |

### The whole genome population structure does not reflect in the effector population structure

The whole genome PCA shows segregation at the continental level; in contrast, the PCA of the annotated effectors shows a complete overlap of the European fields and a partial overlap with the American field (Figure 7. a). The fields MSNs also show an overlap of haplotypes regardless of geographic distribution or between fields variation (Figure 7. a, b; Figure S8. a, b), suggesting a high frequency of sexual reproduction, allowing for every effector haplotype to be present in any field.

**Figure 7.**
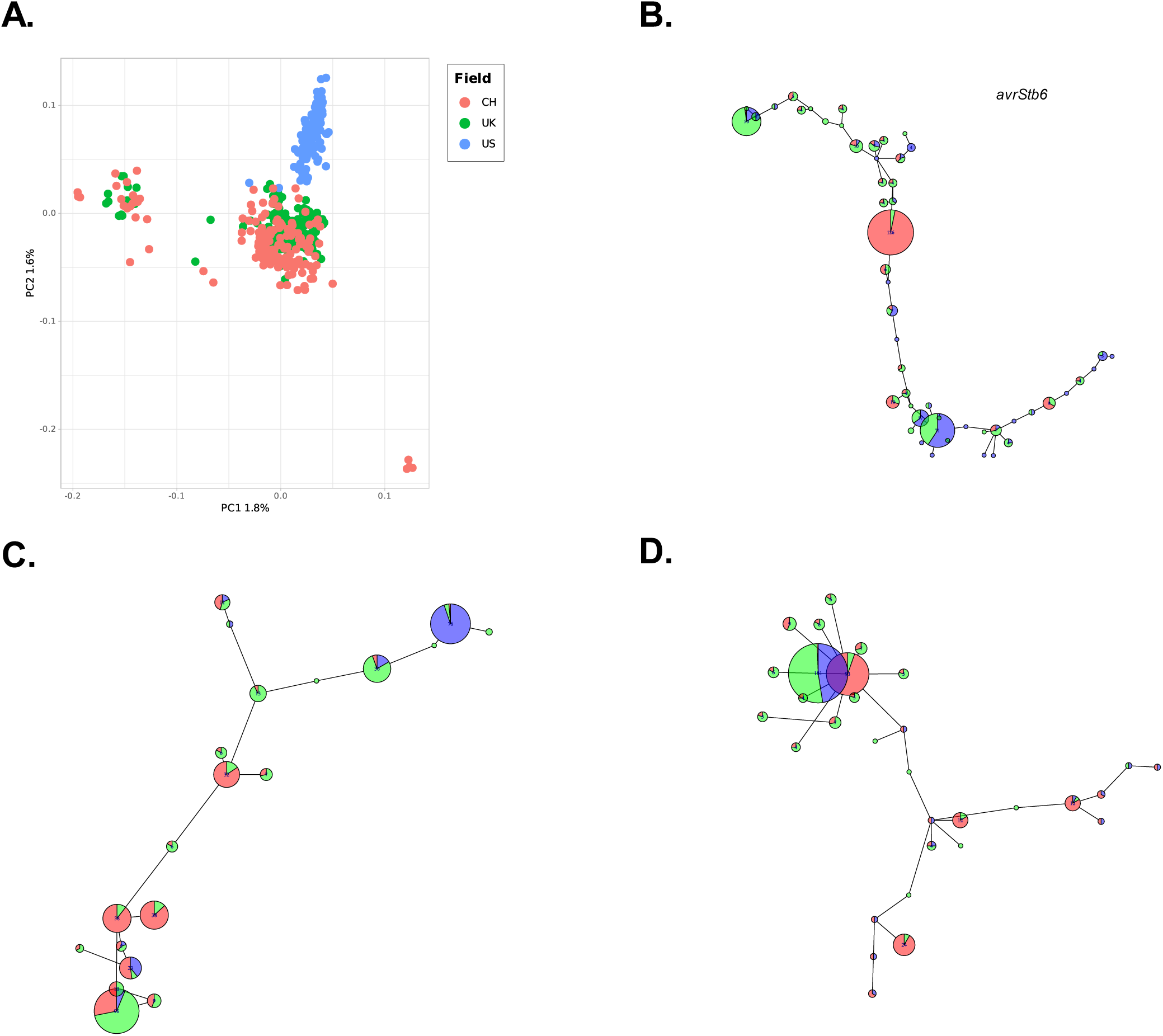
Haplotype distribution for a subset of functionally characterised effectors in the fields. **a.** PCA with the Effectorome (n= 531 genes) SNPs for each field calculated using SNPRelate::snpgdsPCA.. **b.** Minimum spanning network (MSN) for the effector *AvrStb6* between the fields. **c.** MSN for *avr3D1* between the fields. **d.** MSN for *avrStb9* between the fields. Generated using Hamming genetic distance based on SNPs with poppr::bitwise.dist and plotted using poppr::MSN. The Effector annotation and localisation are referenced in Rudd *et al.,* 2015 and Amezrou *et al.,* 2024.

**Figure 8.**
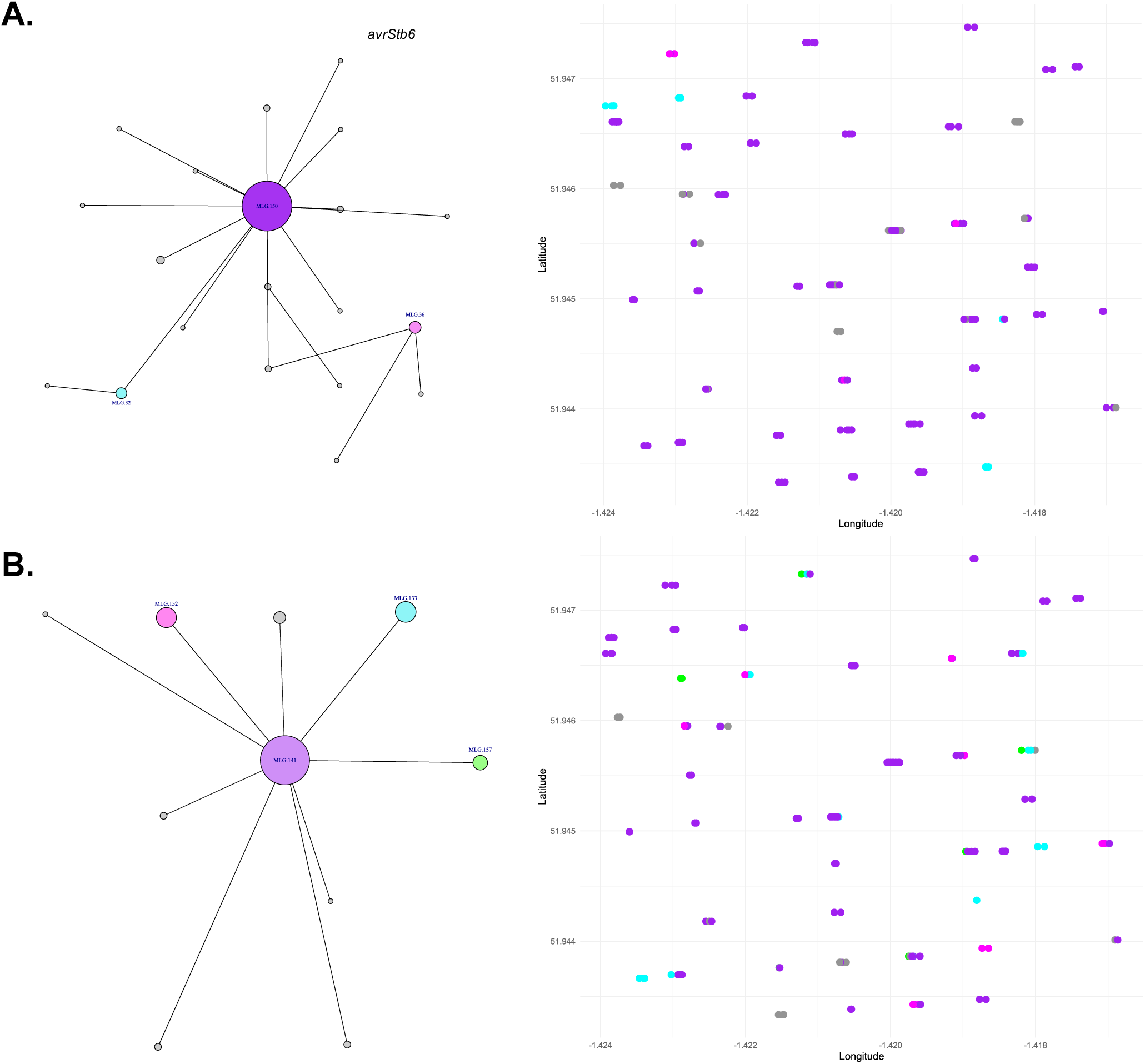
Haplotype geographic distribution for characterised effectors in the UK field. **a.** Minimum spanning network for the effector *avrStb6* and the field’s main Multilocus Genotype geographical distribution. **b.** MSN for *avrStb9* and the main Multilocus Genotype geographical distribution in the field. Generated using Euclidean genetic distance based on SNPs with poppr::bitwise.dist and plotted using poppr::MSN. The Effector genes annotation and localisation were found in Rudd *et al.,* 2015 and Amezrou *et al.,* 2024.

### There is no genomic segregation within the field

The fields’ PCA analyses show no clear genetic structure, with low explanatory power among all principal components. Most of the power is driven by small groups of isolates that are outliers to the condensed cloud of most isolates in the fields (Figure S2b). This confirms that most of the isolates are equally distant from each other, as most SNPs are present in low frequency (Figure 1. a). Nonetheless, for each field, clusters of isolates are more distant from the average. This also suggests that within the fields, there is no clear segregation driven by physical distance, i.e., distance distribution by field lanes or plots. We confirmed this hypothesis in the UK field, where we did a Mantel test for matrix comparisons using the geographic and genetic distance matrix. We show that most pairwise comparisons show a dispersed pattern with no clear relation. The Mantel test was not significant (p < 0.01), which tells us there is no geographic distance-driven separation within the field (Figure S7. b). This is most likely driven by the effect that, in some cases, the isolates taken from a single plant could be genetically different, and a few isolates from different plants clustered together as a clonal cluster but were in different parts of the field (Figure S7. a).

### There is no effector haplotype segregation within the field driven by small-scale geographic distribution

To assess the extent and spread of pathogen effectors within a field, we assess the effectors highlighted before. We calculated haplotype diversity by constructing minimum spanning networks. These show that *avrStb6* has one major haplotype (MLG150) in the field and several rare haplotypes exist, this is also true for avrStb9 (MLG148); in contrast, *ZtIPO323_119610* and *ZtIPO323_117500* show two major haplotypes and then also a few unique isolates in the field (MLG1,2; MLG112, 94 respectively) and avr3D1 shows four major haplotypes (MLG 133,141,152,157) (Figure 7. a, b, Figure S8). Nonetheless, the effector haplotype distribution across the field is also not driven by any pattern related to lanes or similar features. Both effectors could have occurrences of the same haplotype in the same plant or lesion but also have dissimilar haplotypes, which is congruent with the lack of pattern seen in the whole genome (Figure 7. a, b, Figure S9. a, b). In addition to this, we found that in ∼33% of our clonal clusters, we could find a different haplotype for all three effectors.

## Discussion

We first aimed to assess the within-field diversity of *Zt* in three different deeply sampled field sites in Switzerland, the UK and the US. Our comparison with the filtering strategies that have been previously published and our relaxed approach allows us to focus on rare variants and how they impact the differentiation of clonal isolates and the diversity statistics at the field level. Also, it allows for a better understanding of the contribution of dispensable chromosomes to the overall diversity of *Zt*, as the somatic recombination has also been observed in the field before (Hartmann et al., 2017; Möller et al., 2018; Plissonneau et al., 2018; Singh, Karisto, et al., 2021).

With our less stringent approach, we allow an increase in the SNP count of 30% within each field; this does not affect the genetic distance distribution or change factors like clonality. However, it shows a slight change in the observed site frequency spectra, with a minor increase in intermediate-frequency SNPs in all samples. This indicates that we possibly detect putatively adaptive SNPs that have been previously overlooked. The increased SNP count overall allows for a more detailed field characterisation. The newly detected SNPs in intermediate frequencies can become particularly relevant when studying changes over short time frames as different pressures can cause a rapid shift in alleles that would, for example, allow *Zt* to keep overcoming the prophylactic barriers we have placed to control the pathogen, as happened when the Stb16 resistance was broken in France in recent years (Langlands-Perry et al., 2023; Orellana-Torrejon et al., 2022; Suffert et al., 2024) or change to different haplotypes conferring fungicide resistance to new compounds (Birr et al., 2021; Glaab et al., 2024).

Our predictions of the maximum number of SNPs we could encounter in the fields are supported by the particular case of the Swiss field model, where we find that the maximum number of SNPs in the Strict filtering is highly similar to what they reported with 1’497,037 at 177 samples, where our model predicts that the samples needed are 167 with ∼1.5 million SNPs, this shows that the relevant diversity was sufficiently captured with the sampling effort in the field. Our model confirmed the curve’s plateauing with their sampling for relevant variants accumulation threshold of 1% (Singh, Karisto, et al., 2021). The level of the plateau increases to 2 million with our more relaxed settings after ∼170 samples. Yet, we know that the SNPs representing a metapopulation can scale up to more than 20 million following our model fit to the power function. This indetermination of a real plateau suggests that the emergence of different virulent and fungicide resistance haplotypes due to SNPs will remain the biggest challenge in managing STB disease.

With our filtering settings, we found that the European fields have close to 2 million SNPs, and the US field close to 850,000. The high observed within-field diversity in Europe is consistent with the previous studies showing that *Zt* has a very large N_e_ (Croll & McDonald, 2017; Dutta, Hartmann, et al., 2021; Feurtey et al., 2023; Singh, Karisto, et al., 2021). The mixed reproduction strategy maximises the diversity accompanied by the high recombination rate and multiple accessory chromosomes, which have a higher mutation rate, allowing for the presence of non-clonal isolates in a very small distance scale, like what was observed at different leaf layers in the UK (Dutta, Croll, et al., 2021; Karisto et al., 2022; Plissonneau et al., 2018; Singh, Karisto, et al., 2021). The presence-absence variation observed in all the fields is an understudied layer of diversity. These regions could contribute to pathogen aggressiveness as there is evidence of predicted effectors for loci in the dispensable genome and contributions in the timing of the necrotrophic switch of the disease (Lapalu et al., 2023; Plissonneau et al., 2018; Van Westerhoven et al., 2023). The dispensable genome also had more SNPs than the core genome, contributing significantly to the diversity we detected in this study; our less stringent approach allowed us to asses this diversity while keeping the noise of false positive SNPs at a minimum by using the combination of GATK filtering parameters. Still, it kept enough resolution to observe rare events, excluding the failed GATK calls, using the minor allele count set to 1 (Feurtey et al., 2023; Singh, Karisto, et al., 2021). Interestingly, we confirm a short invariable region in chromosome 7 in all the fields containing rDNA that could be due to a recombination event with an accessory chromosome (Schotanus et al., 2015).

The European fields showed a higher SNP abundance than the US. The US also shows a higher nucleotide diversity and Tajima’s D values close to 0, indicating no evidence of selection. Whereas the Swiss and UK fields show low nucleotide diversity and Tajima’s D values close to or below 2, indicating selective sweeps or recent population expansion. In the absence of clear meta-data for the US population, it remains a guess as to whether differences in agricultural practices contribute to this difference. Both European fields were sampled after fungicide treatments, which indeed could cause a bottleneck. The UK and the Swiss fields were samples from a very small area (both less than 400x400m) and from single or multiple susceptible cultivars. The US samples were collected from a highly susceptible (Stephens) and a resistant (Madsen) cultivar. This might allow for the development of more intermediate frequency variation due to the selection of isolates with certain effector haplotypes and lack of sexual recombination during the growing season. Yet, it should be noted that a recent study from France also found relatively low SNP counts of 718,810 in 103 isolates from various fields in France, where the same pattern of low explanatory power of the PCs is observed as in all our assessed fields (Amezrou et al., 2023) (Figure S2). Thus, the total SNP accumulation differences are still an open question for future studies.

Our results show that the genetically distinct groups present at the continental level, where the US separates from the European fields. This is consistent with previous studies (Feurtey et al., 2023). This population structure is probably due to local adaption to the different contexts of management like fungicide regimen, tilling policies, and crop rotation, varietal landscape, among other factors and limited spore transmission across oceans (Birr et al., 2021; Cerón-Bustamante et al., 2023; Hartmann et al., 2021; Haueisen et al., 2019; Vestergård et al., 2023). Interestingly, we could see a similar pattern to what was observed in Feurtey et al., (2023), where the European clade has contributed to the more homogeneous US population. The high potential for SNP accumulation at the field level can also explain why, with reduced sampling like SSR, there is a clear population structure with marked private alleles at the regional level, compared to the whole genome SNP call, the effect of the fitness drivers will be confounded among the mostly neutrally evolving genome (Chedli et al., 2022; Siah et al., 2018).

Next, we specifically focused on within-field effector diversity. Whereas we observe the separation of the three fields in PCA analyses on the whole-genome level, thus separation largely disappears when looking at effectors. Remarkably, *avrStb6*, *avr Stb9, avr3D1, ZtIPO323_117500* or *ZtIPO323_119610* don’t have private haplotypes between the fields, supporting the evidence for high recombination rates and high frequency of sexual reproduction that allows the emergence of multiple haplotypes independently, which are kept in the populations probably due to factor like epiphytic sexual reproduction and multiple host contexts adjacent to one another (Orellana-Torrejon et al., 2022; Orellana-Torrejon et al., 2022). Even at the field scale, in the UK, we see that every possible effector haplotype is randomly distributed in the field, which is evidence of the high impact sexual recombination has during the disease cycle. This dispersed distribution in the field happens for both biotrophic phase-related and necrotrophic phase-related effectors, suggesting that the fitness associated with effector polymorphism has a similar magnitude within the field context (Amezrou et al., 2023; Bernasconi et al., 2022; Hassine et al., 2019; Meile et al., 2018, 2023; Morais et al., 2017; Orellana-Torrejon et al., 2022; Singh et al., 2022; Suffert et al., 2015, 2024). However, we do see that there is more diversity associated with more significantly positive Tajima’s D values in the US compared to the European fields, which corresponds to what happens in the genomic context.

In our clonal clusters in the UK field, we show that multiple effector haplotypes can occur within the cluster, suggesting isolates that have high similarity can still have polymorphism in the Effectorome. This is in line with what was modelled in previous studies, where due to the high number of spores produced during a single disease cycle ( in the order of Trillions), all mutations across the genome have the chance to occur ∼ 1,000 times in *Zt* (B. A. McDonald et al., 2022). The standing variation in *Zt* enables the populations to rapidly break down newly deployed R gene-containing varieties like what happened in France with the ‘Cellule’ cultivar in 2021, this R gene had high success when introduced in 2016, but by 2021 the resistance was completely broken due to changes in the avrStb16q effector (Battache et al., 2022; Saintenac et al., 2021; Suffert et al., 2024). Such effector diversity, even within what we consider clonal isolates, due to the mutation and recombination rate, allowing for an even greater increase of effector haplotype combinations, will challenge traditional management measures like single-gene novel-resistance cultivar deployment or even R gene pyramiding. The causes can be diverse. There is a high probability of multiple effector haplotype configurations arising in a field where the R genes are not deployed. Secondly, R genes are never 100% functional, so some residual, R-gene overcoming, inoculum might slowly accumulate in the assumed resistant cultivar. Thirdly, the mechanism behind R gene-based resistances might be problematic in this highly polycyclic pathogen with long latency: most R genes induce cell death, thus, an R gene response induced by a secondary infection might not just prevent secondary infection but instead, be taken as a cue to the necrotrophic switch, proving inefficient as a resistance mechanism (Bernasconi et al., 2023; Brown et al., 2015; Kettles & Kanyuka, 2016; B. A. McDonald et al., 2022; Stam & McDonald, 2018). Also, effectors like avr3D1, which are highly polymorphic and contribute only to a quantitative response, cause less efficient defence responses against the pathogen and, when in combination with the diverse array of effectors, overcome resistances like Stb7 or Stb12 (Meile et al., 2018, 2023, p. 202). In addition, R genes like stb6 and stb9 also can be partially bypassed by avirulent strains when infiltrated, showing the challenges of quantitative resistance breeding (Battache et al., 2022). Management strategies focused on the reduction of primary infection by means of qualitative resistance, such as R-gene-based approaches, might not prove durable, as it will be impossible to implement through the board deployment of large pyramids of functional R genes and avoid the rise of R-gene overcoming mutations in the tremendously large populations of *Zt.* Instead, integrated solutions focusing quantitatively on symptom reduction and frequent reduction of the inoculum size could prove more durable.

## Acknowledgements

We thank Dr Andreas Buechse for his design for the UK field sampling and ADAS for collecting and preparing the UK field isolates.

