## Supplemental Figure for "*Zymoseptoria tritici* show local differences in within-field diversity and effector variation"

### Supplementary figures for *Zymoseptoria tritici* show local differences in within-field diversity and effector variation

Andrea Tobian Herreno[1], Pu Huang[2], Isabella Siepe[3], Remco Stam[1*]

[1] Institute of Phytopathology Christians-Albrechts University Kiel, Hermann-Rodewald-Strasse 3 9, 24118 Kiel, Germany 4

[2] BASF Corporation, Research Triangle Park, NC 27709-3528, USA 5

[3] BASF SE, APR/FE, Speyerer Strasse 2, 67117 Limburgerhof, Germany 6

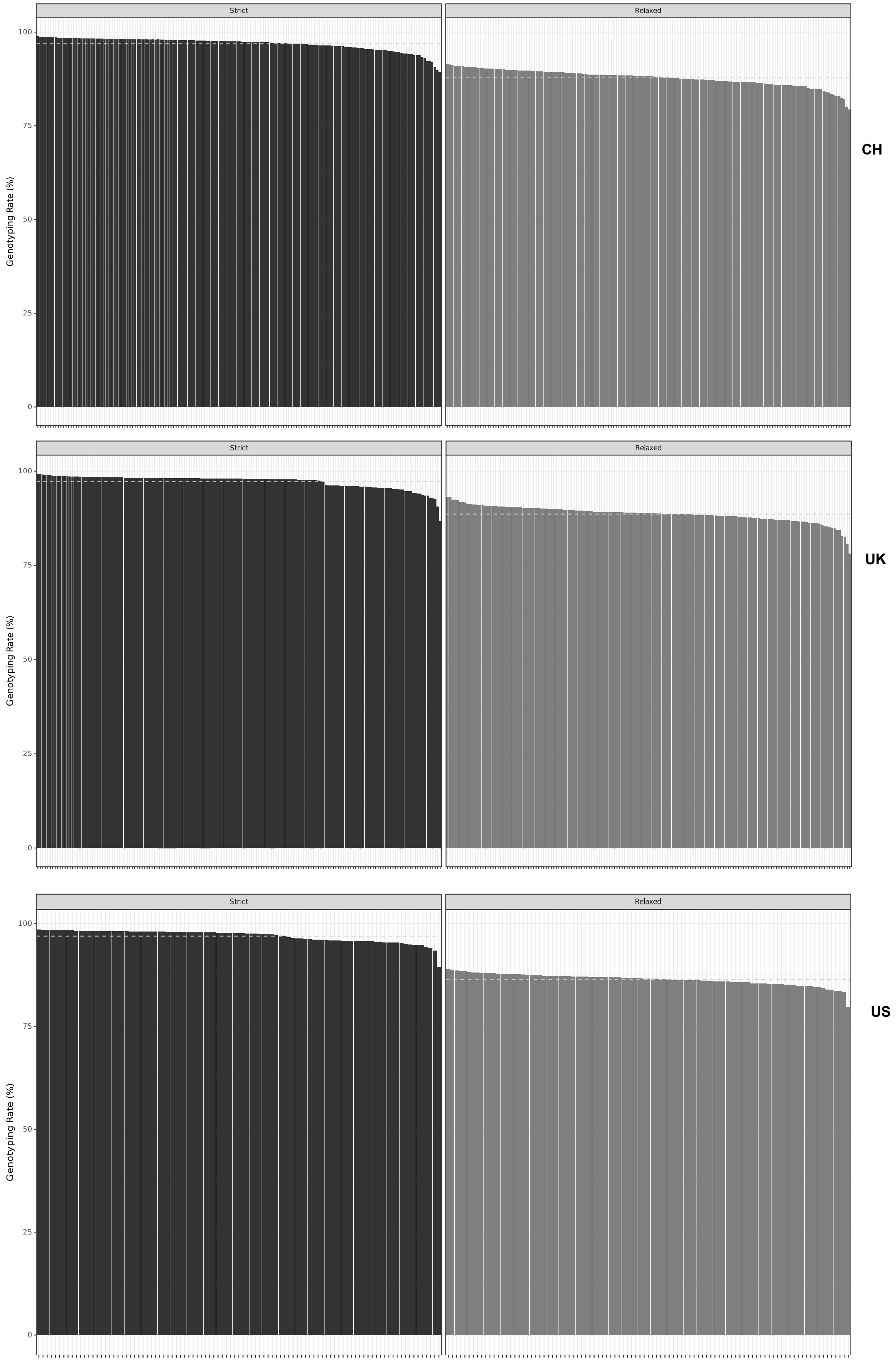


**Figure S1. Genotyping rate of both filtering pipelines and Minor Allele Frequency distribution of the strict-filtered dataset.** The genotyping rate was assessed using PLINK with the --missing option. The Minor Allele Frequency (MAF) distribution was assessed with the -frq option in PLINK. The plots were made using ggplot in R**. a.** Genotyping rate of the Relaxed and Strict filtering. Relaxed filtering dataset has a mean of 87.07% for the UK, 86.04% for CH and 84.25% for the US field. Strict: Genotyping rate of the Singh *et al.* 2021 filtering with a mean of 96.72% for the UK, 97.09% for CH and 96.86% for the US.

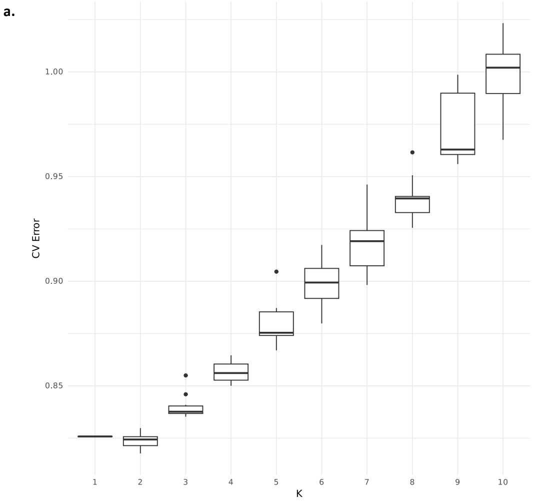


**Figure S2. Cross-validation error for the admixture analysis and Principal Component Analysis of the fields. a.** Cross-validation error scores for the K populations evaluated. K=2 has the lowest CV error and was the most likely scenario. We used Admixture v1.3.0 and used ggplot geom_boxplot() to generate the figure. **b.** PCA from each field shows a repeating pattern of mostly low explanatory power for the core genome, with some isolates driving the explanatory power as outliers. Calculated with SNPRelate::snpgdsPCA ().

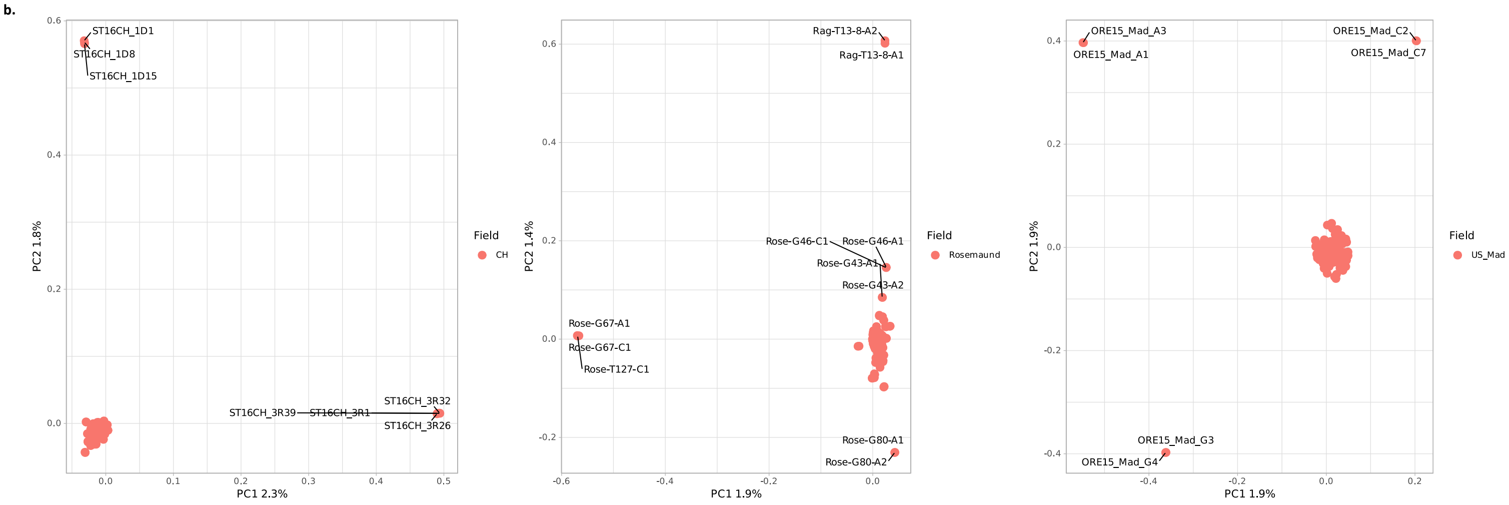

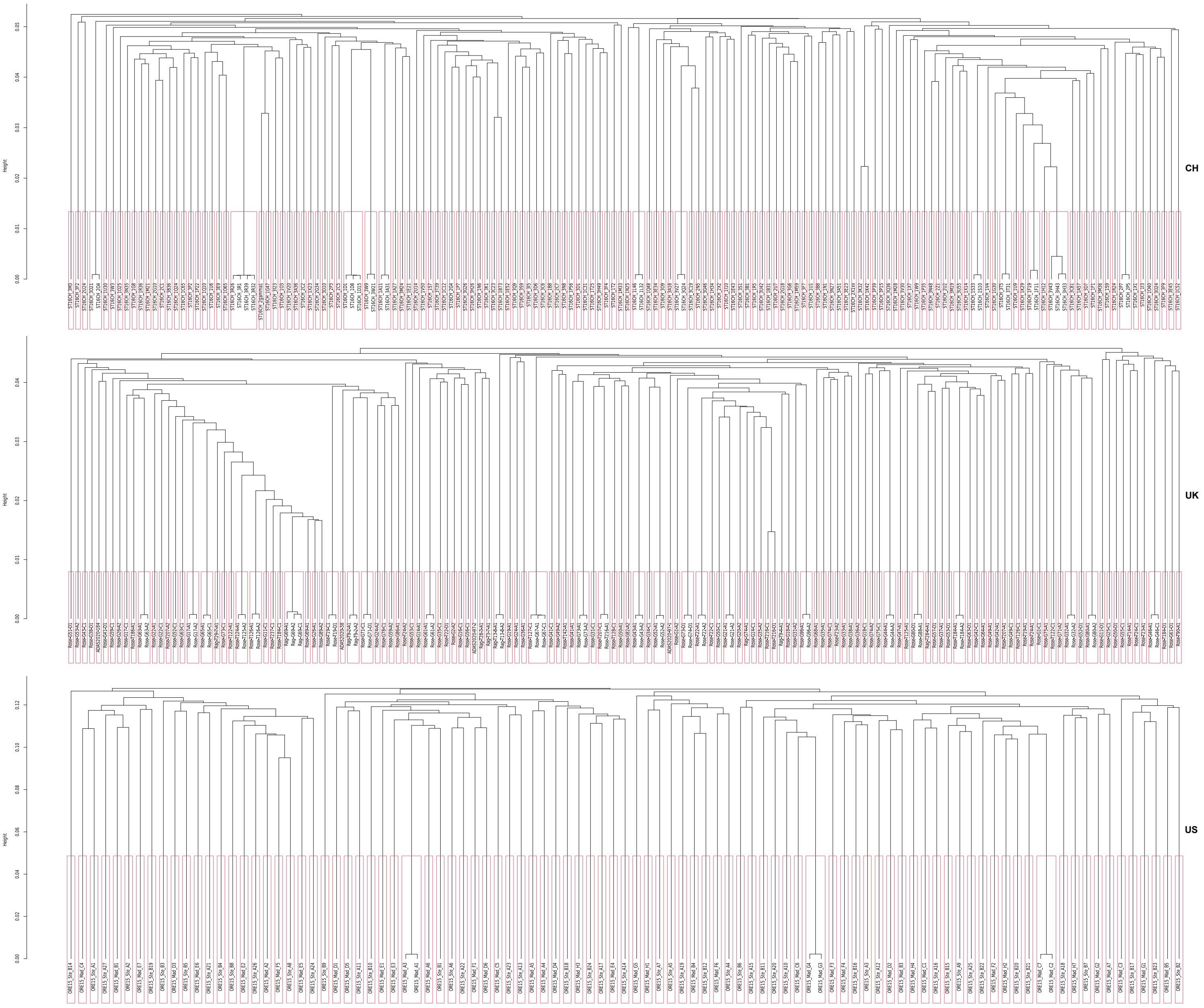


**Figure S3. Neighbour Joining Tree with clonal clusters.** Using the clonal definition of less than 1% dissimilarity, we show the NJT for each field with the clusters generated with the hierarchical clustering R base function hc based on each field's genetic distance matrix made with poppr::bitwise.dist().

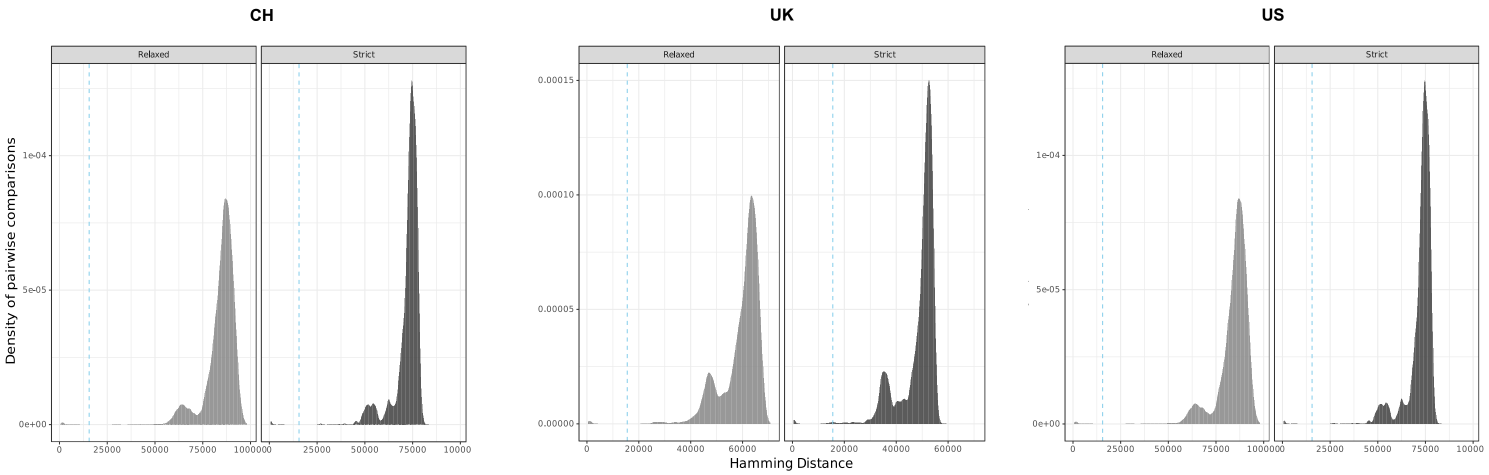


**Figure S4. Genetic Hamming distance disribution with the Strict and Relaxed filters.** CH, UK and US distnace distribution in amount of differing SNPs (Hamming distance) calculated using poppr::bitwise.dist. Sample count was normalised using the ggplot option stat “Density”.

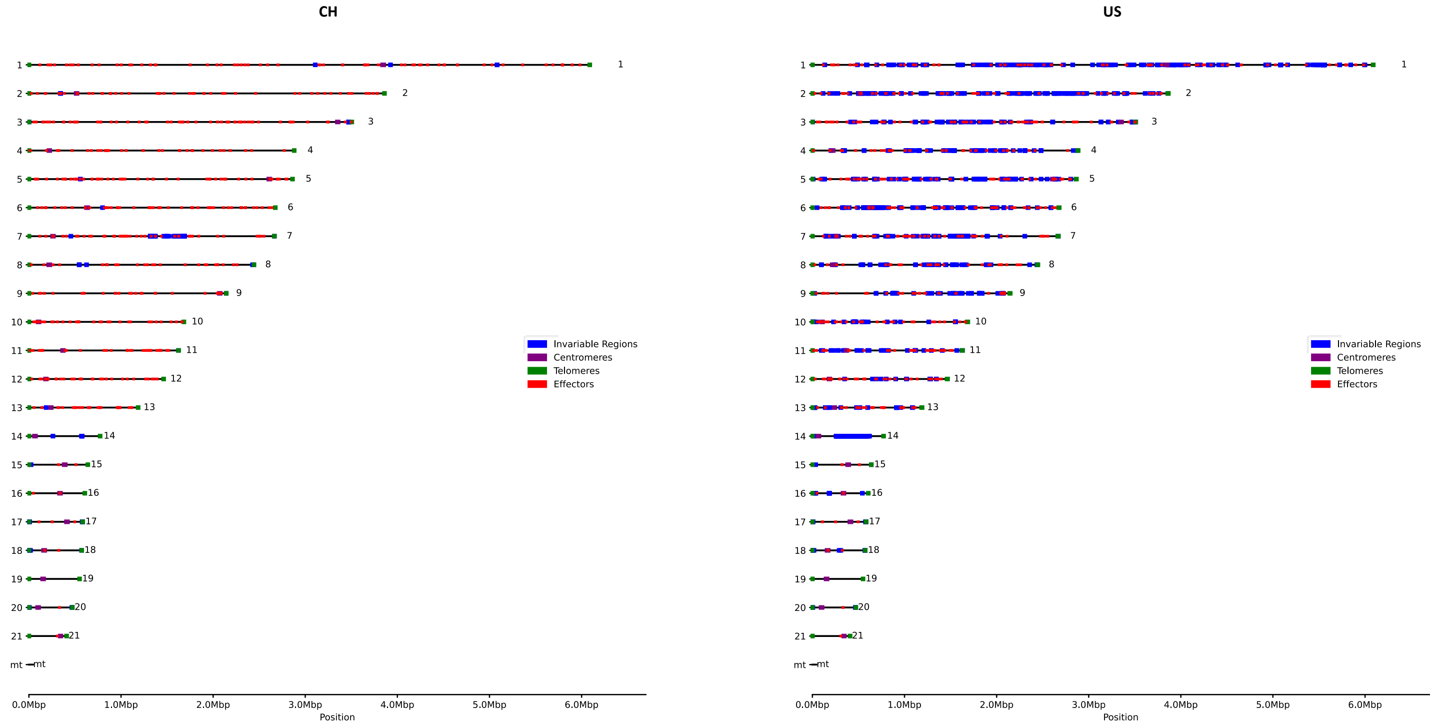


**Figure S5. Invariable regions in the CH and US fields.** The fields were screened for invariable regions per chromosome using the reference IPO323 for comparison. The telomeric and centromeric regions were annotated as determined by Schotanus *et al.* 2015. The invariable regions were corrected using the location for centromeres and telomeres using a custom script.

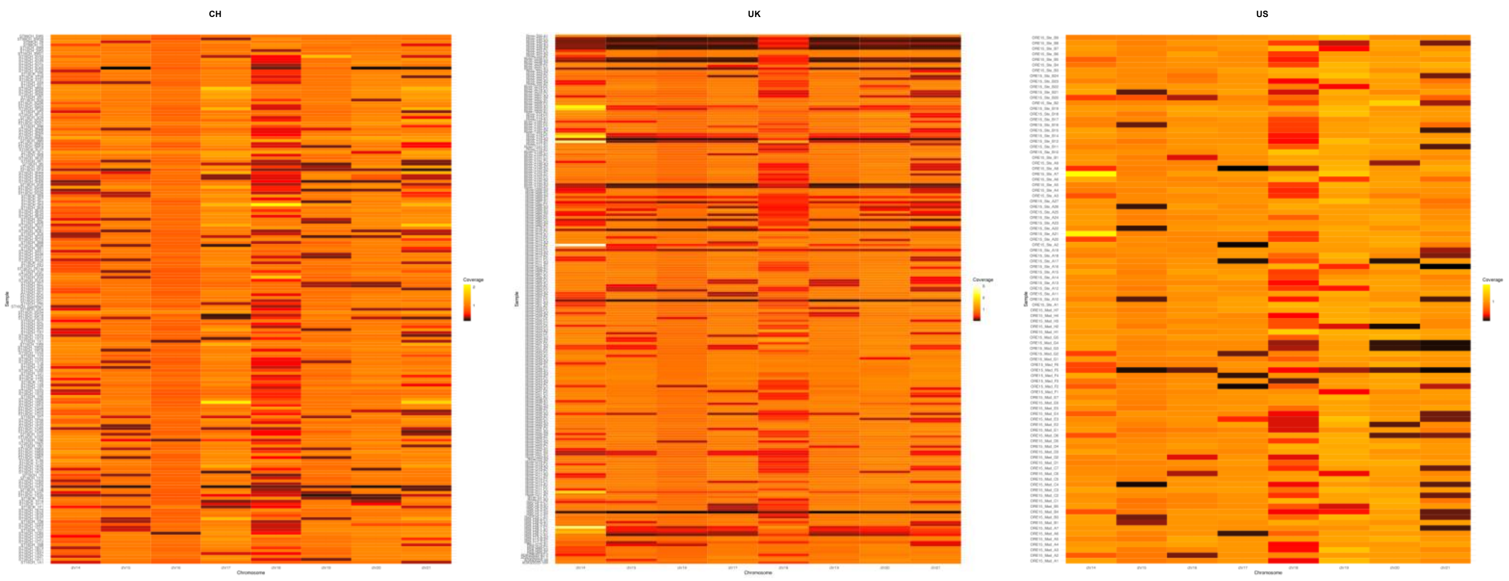


**Figure S6. Accessory chromosome normalised depth.** Depth heatmap normalised by the average depth of the core chromosomes used ggplot geom_tile() to generate the heatmaps of the normalized values and used the following thresholds for the color scale: depth < 0.25 = absent, 0.25 > depth < 0.625 = partial deletion, 0.625 > depth < 1.5 = presence and depth > 1.5 = duplication.

**a.**

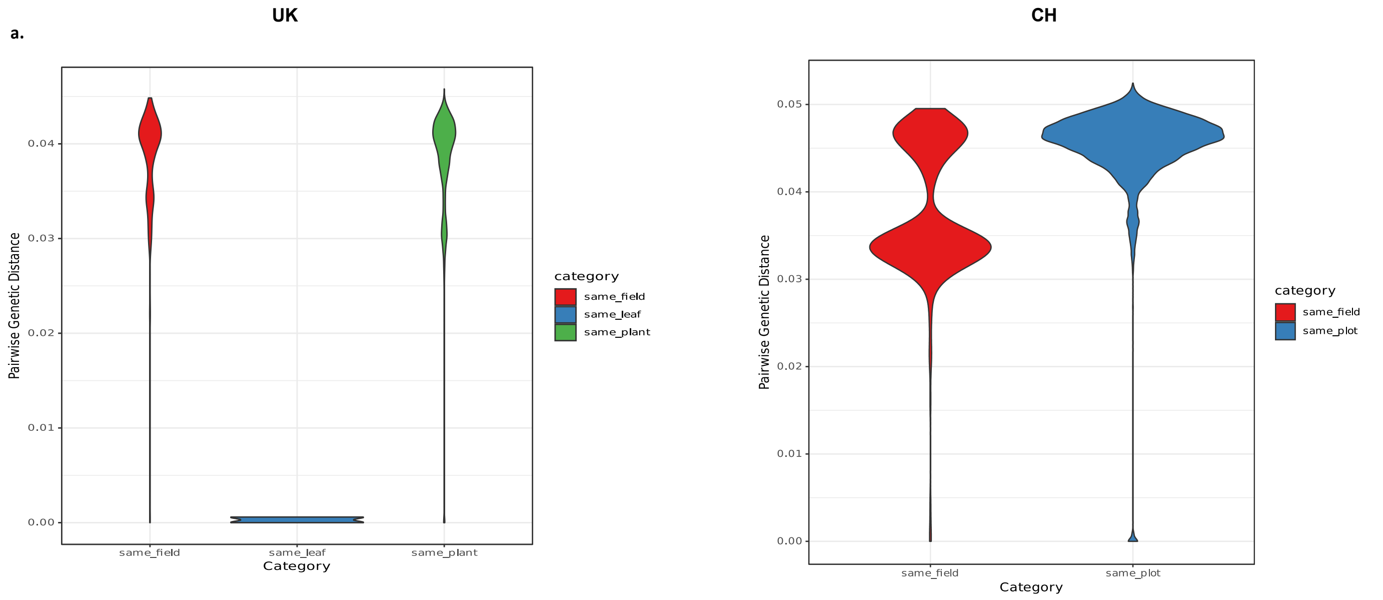


**b.**

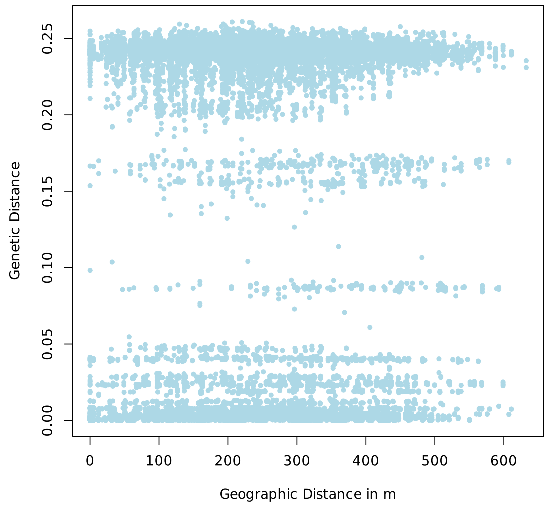


**Figure S7. Genetic distance distribution according to sampling stratification. a.** Violin plots showing the distribution of genetic distances according to collection categories in the available metadata of the fields: sample plant, same leaf and same plot. **b.** Mantel test plot with the comparison of the Pangenome genetic and geographic distance matrixes in the UK field, calculated using geosphere for the geographic distance and poppr::bitwise.dist for the genetic distance.

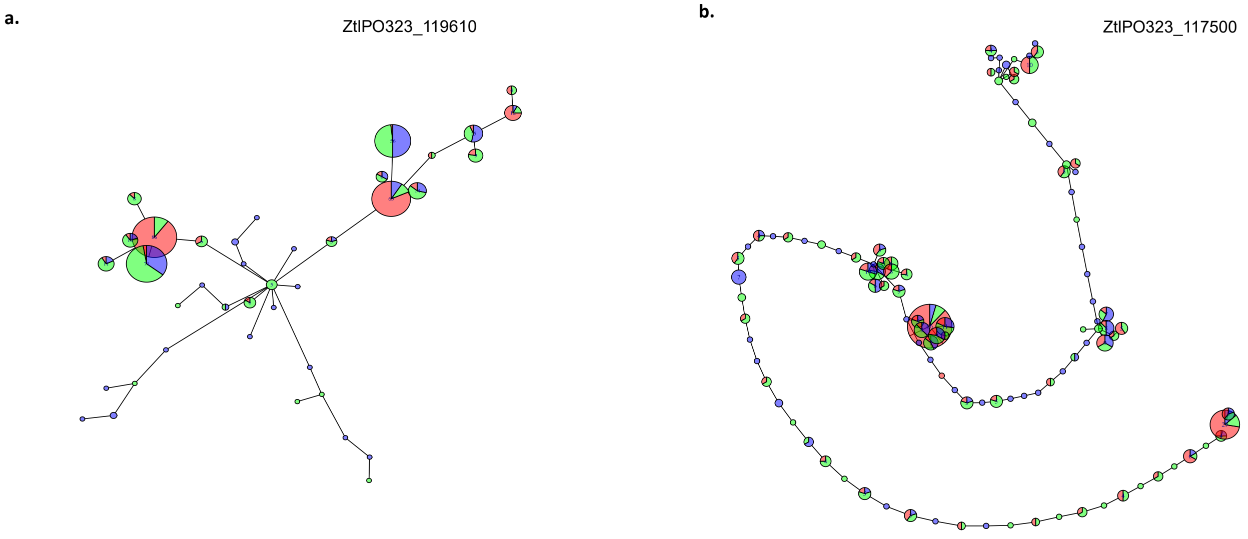


**Figure S8. Haplotype distribution of functionally characterised effectors in the fields. a.** MSN for ZtIPO323_119610 between the fields. **b.** MSN for ZtIPO323_117500 between the fields. Generated using Hamming genetic distance based on SNPs with poppr::bitwise.dist and plotted using poppr::MSN. The Effector annotation and localisation were found in Rudd *et al.,* 2015 and Amezrou *et al.,* 2024.

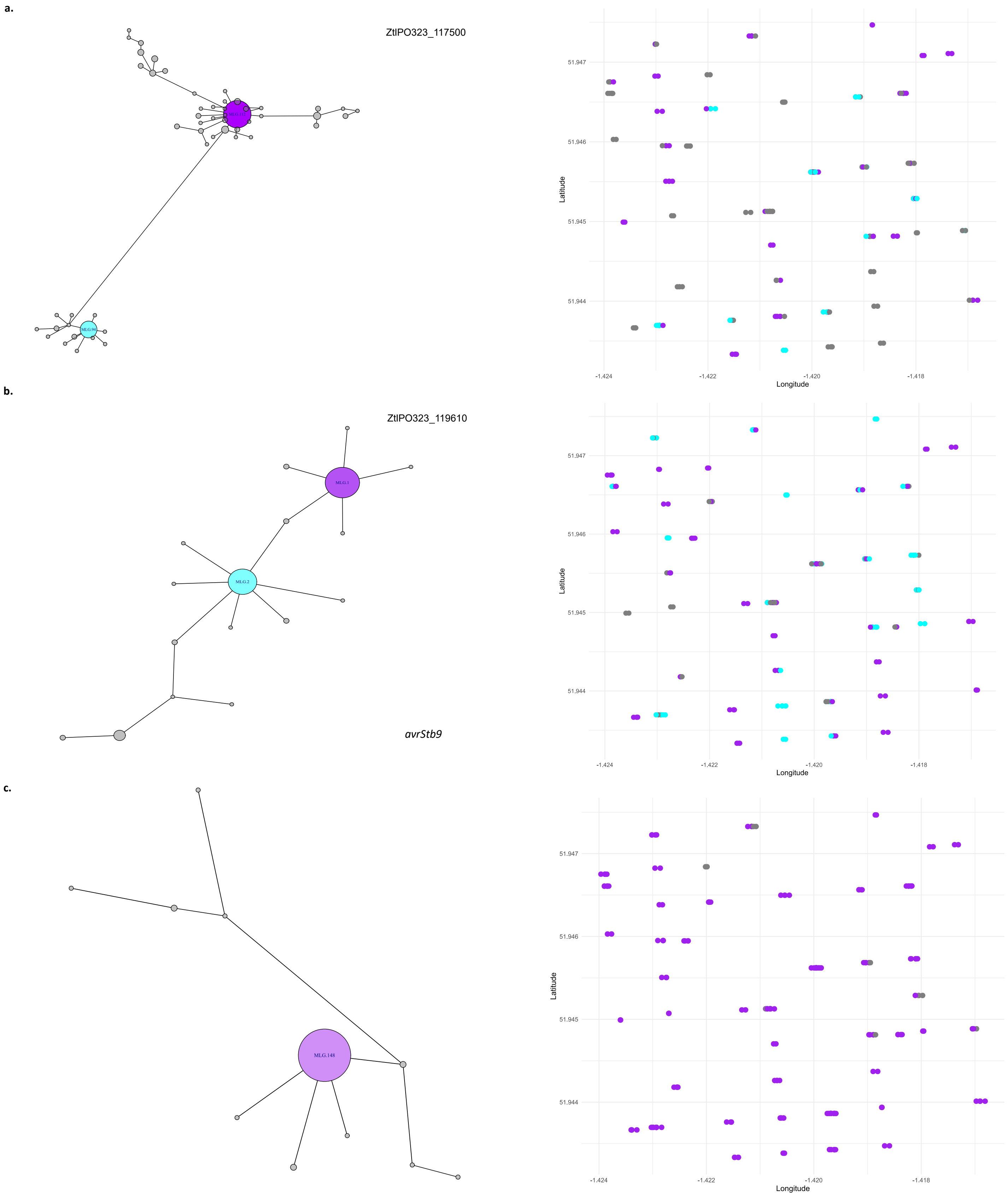


**Figure S9. Haplotype geographic distribution of characterised effectors in the UK field. a.** Minimum spanning network for the effector ZtIPO323_117500 and the field's main Multilocus Genotype geographical distribution. **b.** MSN for ZtIPO323_119610 and the main Multilocus Genotype geographical distribution in the field. **c.** MSN for avrStb9 and the main Multilocus Genotype geographical distribution in the field. Generated using Hamming genetic distance based on SNPs with poppr::bitwise.dist and plotted using poppr::MSN. The Effector genes annotation and localisation were found in Rudd *et al.,* 2015 and Amezrou *et al.,* 2024.
